# Structural snapshots of phenuivirus cap-snatching and transcription

**DOI:** 10.1101/2023.11.29.569195

**Authors:** Harry M. Williams, Sigurdur R. Thorkelsson, Dominik Vogel, Carola Busch, Morlin Milewski, Stephen Cusack, Kay Grünewald, Emmanuelle R.J. Quemin, Maria Rosenthal

## Abstract

Severe fever with thrombocytopenia syndrome virus (SFTSV) is a human pathogen that is now endemic to several East Asian countries. The viral large (L) protein catalyzes viral transcription by stealing host mRNA caps via a process known as cap-snatching. Here, we establish an *in vitro* cap-snatching assay and present three high-quality electron cryo-microscopy (cryo-EM) structures of the SFTSV L protein in biologically relevant, transcription-specific states. In a priming-state structure, we show capped RNA bound to the L protein cap-binding domain (CBD). The L protein conformation in this priming structure is significantly different from published replication-state structures, in particular the N- and C-terminal domains. The capped-RNA is positioned in a way that it can feed directly into the RNA-dependent RNA polymerase (RdRp) ready for elongation. We also captured the L protein in an early-elongation state following primer-incorporation demonstrating that this priming conformation is retained at least in the very early stages of primer extension. This structural data is complemented by *in vitro* biochemical and cell-based assays. Together, these insights further our mechanistic understanding of how SFTSV and other bunyaviruses incorporate stolen host mRNA fragments into their viral transcripts thereby allowing the virus to hijack host cell translation machinery.

## INTRODUCTION

Severe fever with thrombocytopenia syndrome virus (SFTSV) is a segmented negative-strand RNA virus in the order *Bunyavirales*, family *Phenuiviridae* (1). SFTSV has been identified in several East Asian countries, including China, Japan, and South Korea to cause human disease (2–7). Transmitted by ticks, SFTSV is known to infect a range of domestic animals, like cats and dogs, as well as a wide variety of livestock such as cows, chickens, sheep, and pigs (8–11). Although SFTSV is thought to be predominantly transmitted by tick bite, there is also some evidence of human-to-human transmission (12,13). For these reasons, SFTSV is considered a significant threat to global public health with no effective medical countermeasures available.

A key target for the development of new antiviral strategies is the bunyaviral large (L) protein, a multidomain and multifunctional protein essential for both viral genome replication and transcription processes (14). During infection, the negative-sense genomic viral RNA (vRNA) is first copied into a positive-sense complementary RNA (cRNA) intermediate (15). This cRNA is then used as a template to reproduce the negative-sense vRNA, ready for packaging into progeny virions (15). It has been shown for bunyaviruses that viral genome replication can be initiated *de novo* using two different mechanisms: either initiation happens terminally or a short primer is produced internally and subsequently realigned to the 3’ *via* a prime-and-realign mechanism (16–18). For SFTSV, biochemical data has yet to distinguish between the two scenarios; however, recent structural data addressing how the SFTSV L protein enables viral genome replication suggest a prime-and-realign mechanism is more likely (18,19). On the other hand, bunyavirus transcription is thought to be initiated *via* a process known as cap-snatching (20). Similar to influenza virus transcription, short fragments of 5’-capped host cell mRNAs are used by the L protein to prime the synthesis of viral mRNAs (20). This allows the virus to hijack the translation machinery of the infected host cell for the production of viral proteins. Therefore, viral transcription relies on the endonuclease (ENDO) and cap-binding domains (CBD) of the L proteins and the length of the “*stolen*” capped primers appears to be specific to individual bunyaviruses. For example, Lymphocytic Choriomeningitis virus (LCMV, family *Arenaviridae*) uses a relatively short 1 – 7 nt primer (21), whereas Rift Valley fever virus (RVFV, family *Phenuiviridae*) requires a longer primer in the range of 12 – 14 nts (22). Because of their importance in these processes, the isolated cap-binding and ENDO domains of several bunyaviruses have been investigated extensively by X-ray crystallography (18,23–32).

Recently, electron cryo-microscopy (cryo-EM) data from single-particle analysis have become available for several full-length bunyavirus L proteins in complex with viral RNA, including Lassa virus and Machupo virus (LASV and MACV, family *Arenaviridae*), La Crosse virus (LACV, family *Peribunyaviridae*), Sin Nombre Virus and Hantaan virus (SNV and HTNV, family *Hantaviridae*) (16,33–38). This structural data has provided key insights into how these molecular machines interact with the viral RNA promoter and undergo major conformational changes during viral genome replication (39). For SFTSV, high-resolution cryo-EM structures of the L protein in various functionally relevant states of the viral genome replication process, from pre-initiation to late-stage elongation, have also been published (19). This data revealed how the SFTSV L protein binds the 5’ viral RNA in a hook-like conformation and further showed how the 5’ and 3’ RNAs form a distal duplex structure positioning the 3’ RNA terminus in the RdRp active site ready for initiation (19). Further, we could show that at late-stage elongation the L protein core accommodates a 10-bp product-template duplex which splits at the bottom of the RdRp core. In this conformation, the template RNA then binds to a designated 3’ secondary binding site in a similar fashion to that seen for other negative-strand RNA viruses, including LASV and LACV L proteins as well as the influenza virus polymerase complex (19,33,34,40).

These insights notwithstanding, much still remains unknown, especially in relation to how the SFTSV L protein enables viral transcription. Specifically, how does the SFTSV CBD interact with and bind to capped-RNA? How does the L protein present bound capped-RNA fragments to the ENDO for cleavage? How does the L protein then incorporate the cleaved snatched capped-RNA fragment into progeny RNA? Finally, it is still very much an open question how different L protein functions are regulated during infection to switch between *de novo* replication and primed transcription.

To answer these questions, here we used a combination of biochemical *in vitro* assays and single particle cryo-EM to visualise and understand how the SFTSV L protein steals and then incorporates host mRNA fragments into their viral RNA. We establish an *in vitro* cap-snatching assay and present 3 cryo-EM structures of the SFTSV L protein captured during viral transcription. The data are refined to an overall resolution of 2.98 – 3.40 Å. This structural data is validated by a comprehensive mutational analysis in a cell-based mini-replicon system. Together, these insights further our understanding of how SFTSV and other bunyaviruses steal and incorporate host mRNA fragments into their viral RNA thereby allowing the virus to hijack the host cell translation machinery.

## MATERIALS AND METHODS

### Expression and purification of SFTSV L protein

The L gene of SFTSV AH 12 (accession no. HQ116417) containing a C-terminal StrepII-tag was chemically synthesised (Centic Biotech, Germany) and cloned into an altered pFastBacHT B vector (18). Point mutations (D112A) were introduced by mutagenic PCR to reduce the SFTSV L proteins intrinsic endonuclease activity (31). DH10EMBacY *E. coli* cells were used to generate recombinant baculoviruses, which were subsequently used for L protein expression in Hi5 insect cells. The harvested Hi5 insect cells were resuspended in Buffer A (50 mM HEPES[NaOH] pH 7.0, 1 M NaCl, 10% [w/v] Glycerol and 2 mM DTT), supplemented with 0.05% (v/v) Tween20 and protease inhibitors (cOmplete mini, Roche), lysed by sonication, and centrifuged twice (20,000 x g for 30 minutes at 4°C). The soluble protein fraction was loaded on Strep-TactinXT beads (IBA Lifesciences) and eluted with 50 mM Biotin (Applichem) in Buffer B (50 mM HEPES[NaOH] pH 7.0, 500 mM NaCl, 10% [w/v] Glycerol and 2 mM DTT). L protein-containing fractions were pooled and diluted 1:1 with Buffer C (20 mM HEPES[NaOH] pH 7.0) before loading on a heparin column (HiTrap Heparin HP, Cytiva). Proteins were eluted with Buffer A and concentrated using centrifugal filter units (Amicon Ultra, 30 kDa MWCO). L proteins then used for single-particle cryo-EM were further purified by size-exclusion chromatography using a Superdex 200 column (Cytiva, formerly GE Life Sciences) in buffer B. Purified L proteins were concentrated as described above, flash frozen in liquid nitrogen and stored at −80°C.

### Polymerase assay

#### RNA preparation

Primer A (1–24) and B (1–32) were chemically synthesized with a 5’ diphosphate modification (Chemgenes) (Supplementary Table 1). An N7-MeGppp (cap^0^) was introduced at the 5′ terminus of Primer A (1–24) and B (1–32) using the ScriptCap m^7^G Capping System (CELLSCRIPT) with 50 μg oligo, according to the manufacturer’s standard protocol. After the addition of cold AmOAc (2.5 M) the capped RNA was precipitated overnight in EtOH (>99%) at −20°C, washed two times with EtOH (80%), dried and dissolved in DEPC treated H_2_O at room temperature. The capped RNA was purified further by passing through a MicroSpin G-25 column, as per the manufacturer’s instructions.

#### In vitro cap-snatching assay

For the *in vitro* cap-snatching assay, 0.8 μM of the wild-type SFTSV L protein was incubated sequentially with single-stranded L 5’ (1–20) RNA and single-stranded L 3’-6A (1–26) modified with 6 additional A bases, at a x1.1 molar excess to the L protein in assay buffer (100 mM HEPES, 50 mM KCl, and 2 mM DTT, pH 7) on ice for 5 minutes (Supplementary Table 1). To this, either Primer A (1–24) or B (1–32) was added to a final concentration of 4.5 μM and the reaction incubated on ice for additional 30 minutes. The reaction was started by the addition of NTPs (0.2 mM ATP/UTP/GTP, 0.1 mM CTP spiked with 166 nM 5 µCi[α]32P-CTP) and MgCl_2_ (final concentration 2.5 mM) in a final reaction volume of 10 μL. After incubation at 30°C for 1-60 minutes, the reaction was stopped by adding an equivalent volume of RNA loading buffer (98% formamide, 18 mM EDTA, 0.025 mM SDS, xylene cyanol and bromophenol blue) and heating the samples at 95°C for 5 minutes. Products were then separated by denaturing gel electrophoresis using 25% polyacrylamide 7 M urea gels and 0.5-fold Tris-borate-EDTA running buffer. Signals were visualised by phosphor screen autoradiography using a Typhoon FLA-7000 phosphorimager (Fujifilm) operated with the FLA-7000 software.

### Rift valley fever virus mini-replicon system

The experiments were performed using the RVFV mini-replicon system described previously (41). L genes were amplified using mutagenic PCR from a pCITE2a-L template to produce either wild-type or mutated L gene expression cassettes. PCR products were gel purified and quantified spectrophotometrically. All mutations were confirmed by sequencing of the PCR products. BSR-T7 cells were transfected per well of a 24 well plate with 250 ng of L gene PCR product, 500 ng of pCITE expressing NP, 750 ng of pCITE expressing the mini-genome RNA encoding Renilla luciferase (Ren-Luc), and 10 ng of pCITE expressing the firefly luciferase as an internal control. At 24 hours post-transfection, either total cellular RNA was extracted for Northern blot analysis using an RNeasy Mini kit (Qiagen) or cells were lysed in 100 μL of passive lysis buffer (Promega) per well, and firefly luciferase and Ren-Luc activity quantified using the dual-luciferase reporter assay system (Promega). Ren-Luc levels were corrected with the firefly luciferase levels (resulting in standardised relative light units [sRLU]) to compensate for differences in transfection efficiency or cell density. Data were evaluated in Prism and are always presented as the mean of a given amount (n) of biological replicates as well as the respective standard deviation (SD).

### Northern blot analysis

For the Northern blot analysis, 600-750 ng of total cellular RNA was separated in a 1.5% agarose-formaldehyde gel and transferred onto a Roti-Nylon plus membrane (pore size 0.45 μΜ, Carl Roth). After UV cross-linking and methylene blue staining to visualise 28 S rRNA, the blots were hybridised with a 32P-labelled riboprobe targeting the Ren-Luc gene. Transcripts of the Ren-Luc gene and complementary replication intermediate RNA of the minigenome were visualised by autoradiography using a Typhoon FLA-7000 phosphorimager (Fujifilm) operated with the FLA-7000 software. Quantification of signals for antigenomic RNA and mRNA was done in Fiji. To confirm general expressability of the L protein mutants in BSR-T7 cells, the cells were transfected with 500 ng L gene PCR product expressing C-terminally 3xFLAG-tagged L protein mutants per well. Cells were additionally infected with Modified Vaccinia virus Ankara expressing a T7 RNA polymerase (MVA-T7) to boost the expression levels and, thereby facilitate detection by immunoblotting. At 24 hours post-transfection, cells were lysed in 50 μL passive lysis buffer (Promega) per well. After cell lysis and separation in a 3-8% Tris-acetate polyacrylamide gel (Invitrogen), proteins were transferred to a nitrocellulose membrane (Cytiva). FLAG-tagged L protein mutants were detected using horseradish peroxidase-conjugated anti-FLAG M2 antibody (1:9,000) (A8592; Sigma-Aldrich) and bands were visualised by chemiluminescence using Clarity Max Western ECL Substrate (Bio-Rad) and a FUSION SL image acquisition system (Vilber Lourmat).

### Sample preparation for cryo-EM

#### TRANSCRIPTION-PRIMING and TRANSCRIPTION-PRIMING (in vitro) structures

The SFTSV L protein at a concentration of 3 μΜ in assay buffer (100 mM HEPES, 50 mM KCl, 2.5 mM MgCl2, and 2 mM DTT, pH 7) was mixed sequentially with single-stranded L 5’ (1–20) RNA and single-stranded L 3’-6A (1–26) RNA modified with 6 additional A bases, in a x6 molar excess. Single-stranded capped-RNA (Primer E, 1-17, Supplementary Table 1) was then added and the L protein-RNA mixture incubated on ice for 30 minutes. After incubation at 30°C for 5 minutes, the sample was then centrifuged at 15000 x g for 10 minutes at 4°C. Aliquots of 3 μL were applied to Quantifoil R 2/1 Au G200F4 grids, immediately blotted for 2 seconds, and plunge frozen into liquid ethane/propane cooled to liquid nitrogen temperature using a FEI Vitrobot Mark IV (4°C, 100% humidity, blotting force −10).

#### TRANSCRIPTION-EARLY-ELONGATION structure

The SFTSV L protein at a concentration of 3 μΜ in assay buffer (100 mM HEPES, 50 mM KCl, 2.5 mM MgCl2, and 2 mM DTT, pH 7) was mixed sequentially with single-stranded L 5’ (1–20) RNA and single-stranded L 3’-mid AAA (1–20) RNA where nts 6 – 8 have been replaced by A’s, in a x1.1 molar excess. Single-stranded capped-RNA (Primer E, 1-17, Supplementary Table 1) was then added and the L protein-RNA mixture incubated on ice for 30 minutes, after which the reaction was initiated by addition of NTPs (0.63 mM ATP/CTP/GTP/NhUTP). After incubation at 30°C for 2 hours, the sample was then centrifuged at 15000 x g for 10 minutes at 4°C. Aliquots of 3 μL were applied to Quantifoil R 2/1 Au G200F4 grids, immediately blotted for 2 seconds, and plunge frozen into liquid ethane/propane cooled to liquid nitrogen temperature using a FEI Vitrobot Mark IV (4°C, 100% humidity, blotting force −10).

### Cryo-EM single-particle analysis

#### Data collection

Grids were loaded into a 300-kV Titan Krios transmission electron microscope (Thermo Fisher Scientific) equipped with a K3 direct electron detector and a post-column GIF BioQuantum energy filter (Gatan). Micrograph movies were typically collected using the EPU software (Thermo Fisher Scientific) at a nominal magnification of either 105000x with a pixel size of 0.85 Å using a defocus range of –0.8 to –2.6 μm (TRANSCRIPTION-PRIMING and TRANSCRIPTION-PRIMING (*in vitro*) structures) or a nominal magnification of 130 000x with a pixel size of 0.67 Å using a defocus range of –0.8 to –2.4 μm (TRANSCRIPTION-EARLY-ELONGATION structure). The samples were exposed to a total dose of 50 electrons/Å^2^. Recorded movie frames were fractionated into 50 frames. A summary of the key data collection statistics for the dataset used to determine each cryo-EM structure is provided in Supplementary Table 2.

#### Image processing

All movie frames were imported in RELION 4.0 (42) and motion corrected using its own implementation of MotionCor2 (43). Particles were picked automatically using the pre-trained BoxNet2Mask_20180918 model in Warp (44). Calculation of contrast transfer function (CTF) estimation and correction was done in RELION 4.0 with CTFFIND4 (45). Picked particles were extracted and processed in RELION 4.0. First, particles were subjected to 2D classification at 4x binned pixel size. Then, all non-junk particles were used for 3D classification in an iterative manner using a 40 Å low pass filtered map of the apo SFTSV L protein (PDB: 6Y6K) (18). Incomplete, low resolution or 3D classes containing damaged particles were excluded from further data analyses. The remaining particles were re-extracted with the original pixel size value for further 3D classification and refinement. The processing pipeline for each structure determination is given in Supplementary Figure 1 (TRANSCRIPTION-PRIMING and TRANSCRIPTION-PRIMING (*in vitro*) structures) and Supplementary Figure 2 (TRANSCRIPTION-EARLY-ELONGATION structure). All cryo-EM maps were CTF-refined and Bayesian-polished before final refinement in RELION 4.0. All final cryo-EM density maps were generated by post-processing in RELION 4.0. The resolutions of the cryo-EM density maps were estimated using the 0.143 gold standard Fourier Shell Correlation (FSC) cut-off. A summary of the key processing statistics for each cryo-EM structure determination is provided in Supplementary Table 2.

#### Model building, refinement and validation

The recently published SFTSV EARLY-ELONGATION structure (PDB: 8AS7) was used as the starting point for modelling of the structures published in this study (19). These initial models underwent iterative rounds of model building in Coot (46) followed by real-space refinement in Phenix (47). The comprehensive validation tool in Phenix was then used to assess model quality (47,48). Coordinates and EM maps are deposited in the PDB and EMDB databases, as indicated in Supplementary Table 2. Figures displaying the molecules have been generated using a combination of CCP4mg (49), Pymol (Schrödinger), and UCSF ChimeraX (50).

## RESULTS

Here, we aimed to functionally and structurally characterise the transcription-related activities of the SFTSV L protein as a model phenuivirus. To this end, we established an *in vitro* cap-snatching reaction in which we can visualise capped RNA cleavage and subsequent incorporation of the capped primer into transcription products. We also report three high-resolution structures of the SFTSV L protein representing snapshots of the viral genome transcription process by single-particle cryo-EM providing a structural basis for phenuivirus cap-snatching. An overview of the structures including their key features is provided in Table 1.

**Table 1.** Overview of Cryo-EM Structures. An overview of the cryo-EM structures is provided with the reported resolution (Å), any RNA ligands present, PDB and EMDB accession codes, as well as supporting comments highlighting the key features of each structure.

| Name | Res (Å) | RNA | PDB Code | EMDB Code | Comment |
| --- | --- | --- | --- | --- | --- |
| **TRANSCRIPTION-PRIMING** | 3.35 | L 5' (1-20)<br>L 3'-6A (1-26)<br>Primer E (1-17) |  |  | The L protein N- and C-terminal domains are largely rearranged enabling the CBD to feed the bound capped-RNA into the RdRp core. The L protein is also bound to the 5' and 3' RNAs, which form a distal duplex structure, with the 3' RNA feeding into the RdRp core. The tail-end of the capped RNA is shown to base-pair with the 3' RNAs. Although the 5' and 3' ends of the capped primer can be fit with some confidence, it is not possible to fit the entire length of the primer into the map. |
| **TRANSCRIPTION-EARLY-ELONGATION** | 3.40 | L 5' (1-20)<br>L 3'-mid AAA (1-20)<br>Primer E (1-17) |  |  | The L protein N- and C-terminal domains remain in the priming state. The L protein is bound to the 5' and 3' RNAs, which form a distal duplex structure and the 3' RNA feeds into the RdRp core. However, the 3' RNA between the distal duplex and the entrance to the RdRp core appears under strain/tension. There is a stalling |
|  |  |  |  |  | NhUTP at the +1 position indicating that elongation has started but not progressed far. |
| **TRANSCRIPTION-PRIMING (*in vitro*)** | 2.98 | L 5' (1-20)<br>L 3'-6A (1-26)<br>Primer E (1-17) |  |  | The L protein N- and C-terminal domains are found in their 'apo' positions with the ENDO now sitting across the floor of the RdRp core. The L protein is bound to the 5' and 3' RNAs, which form a distal duplex structure, with the 3' RNA feeding into the RdRp core. There are nts shown base-pairing with the 3' RNA in the RdRp core, however, given NTPs were missing for this reaction, these nts are likely primer. The capping reaction is not 100% effective and so we have a mixture of triphosphorylated primer and capped primer in the reaction mix. The primer base-pairing with the 3' RNA in the RdRp core here is likely therefore uncapped primer, hence, explaining why the CBD is not involved in comparison with the TRANSCRIPTION-PRIMING structure. |

### Establishing an *in vitro* cap-snatching reaction

To explore cap-snatching *in vitro*, we incubated the wild-type SFTSV L with three RNA species: 5’ (1–20) and 3’-6A (1–26) RNA, which correspond to the SFTSV genome termini (large segment), and a 32 nucleotides (nt) primer modified with the addition of either a diphosphate or cap^0^ at the 5’ end making for a total length of 33 nts (Primer C, Supplementary Table 1). The rationale behind the length of this primer stems from sequencing data showing that phenuiviruses typically snatch primers in the range of 12-14 nts (22). Thus, we wanted to ensure the primer was sufficiently long so as to engage both the CBD and ENDO. The reaction mix was incubated at 30°C between 1 and 60 minutes with samples taken at regular time intervals. In the presence of an uncapped diphosphorylated primer, the L protein produces a major product at around 52-54 nts (Figure 1). Here, the primer is at a 5-fold excess to the 5’ and 3’ RNAs and it is noteworthy that we do not see any clear evidence of a ‘*de novo*’ replication product at the expected size of 26 nts. We propose that the major product at 52-54 nts corresponds to the L protein using the full-length primer to initiate RNA synthesis. This can be achieved *via* the 5’ end of the primer, which consists exclusively of AC repeats (n=10), base-pairing with the terminal 3’-UGUG…-5’ of the 3’ RNA. This product band is noticeably smeared, which would indicate that the primer and template are base-pairing in a variable fashion rather than consistently (Figure 1). For example, the terminal AC repeat on the primer could base-pair with the terminal UG of the 3’ RNA or, alternatively, the terminal ACAC on the primer could base-pair with the terminal UGUG of the 3’ RNA. This would then lead to similarly-sized products that vary by 2 nts. Formation of this major 52-54 nts product is clearly detectable from the 5 minute timepoint onwards and then gradually decreases in intensity until the 1 hour timepoint (Figure 1), when the band is very faint. This is accompanied by a clear shift in the band profile at the lower end of the gel, where the intensity of the smaller ‘*product’* bands increases with time (Figure 1). These bands most likely correspond to fragments of RNA that have been degraded over time by the SFTSV L protein endonuclease.

**Figure 1.**
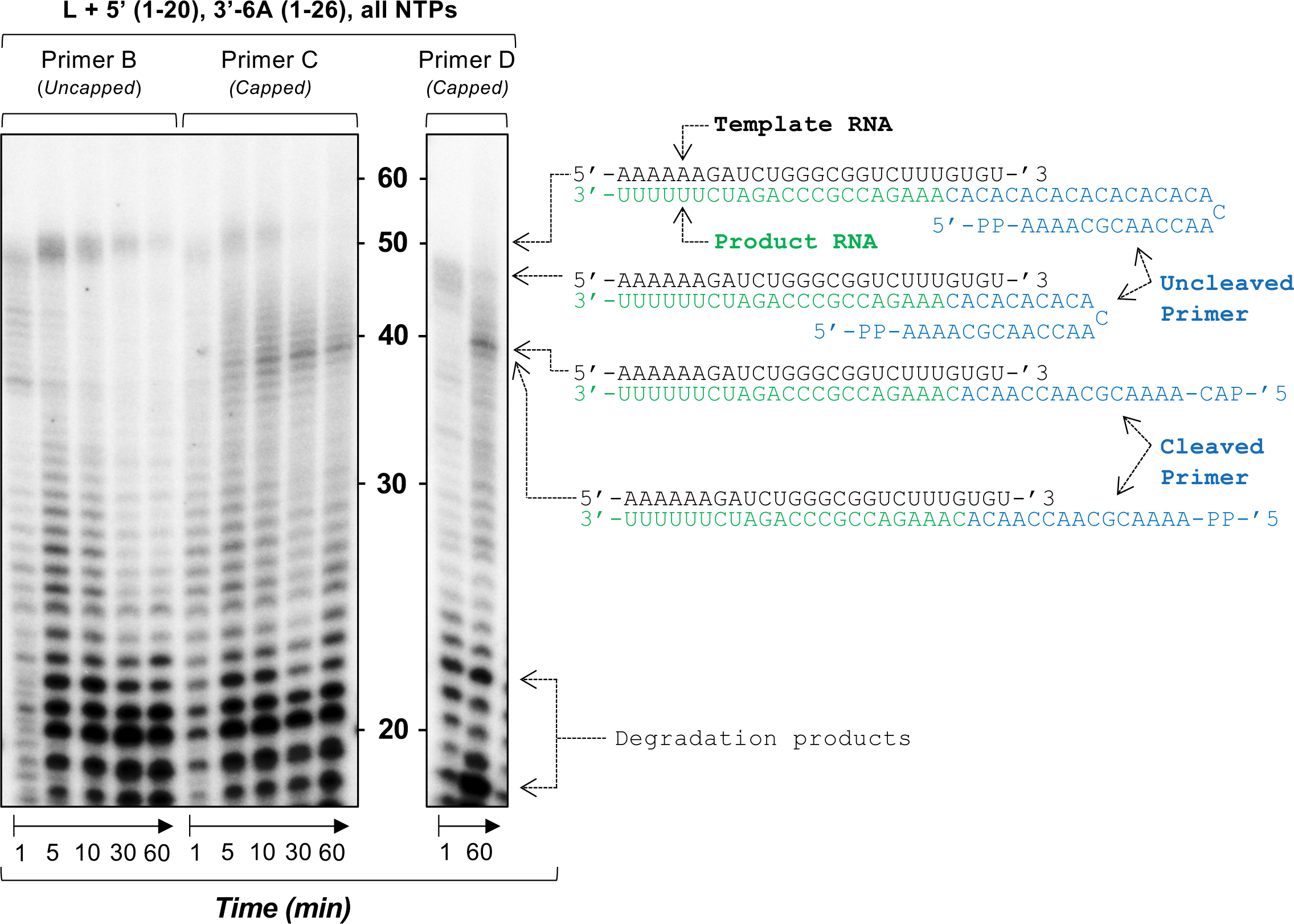
Establishing an *in vitro* cap-snatching reaction. (Left Panel) The impact of primer capping in the presence of the wild type SFTSV L protein was tested *in vitro*. The reactions were carried out with the L 5’ (1–20), L 3’-6A (1–26), and a 32 nt primer that was either diphosphorylated at the 5’ end or capped through the addition of an m7GTP cap under standard polymerase assay conditions (see Materials and Methods). (Right Panel) The impact of primer length was then tested *in vitro*. The reactions were carried out with the wild type L protein, which was incubated with the L 5’ (1–20), L 3’-6A (1–26), and a 24 nt primer that was capped through the addition of an m7GTP cap at the 5’ end as per our standard polymerase assay conditions (see Materials and Methods). For both reactions, aliquots were taken at regular time intervals from 1 minute to 60 minutes. Products were separated by denaturing gel electrophoresis and visualized by autoradiography.

In the presence of the 32-mer capped RNA, the major product appears to be at 39 nts rather than 52-54 nts (Figure 1). We propose this corresponds to a cleaved-capped primer-incorporated product wherein the primer was generated by cap-snatching: the capped RNA is bound initially by the CBD, then cleaved around nt 15 by the ENDO and finally inserted into the RdRp active site. Here, the primer base-pairs with the template enabling RNA synthesis initiation. This primer length would fit with the reported 12-14 nts of non-viral sequences at the 5’ ends of other phenuivirus transcripts (Figure 1) (22). The capped RNA fragment produced would end at the 3’ terminus with an ACA which could then base-pair with the terminal UGU of the 3’ template RNA. In a similar fashion to the uncapped primer reaction, we see noticeable amounts of cleaved capped-primer incorporated product from the 5-10 minute timepoint with endonuclease-mediated degradation then apparent as the reaction proceeds (Figure 1). Curiously, there remains a small amount of full-length product in which the primer apparently has not been cleaved (Figure 1). We think the reason for this is that the capping reaction is not 100% efficient, therefore we expect that in our capped RNA preparation, a small amount of uncapped disphosphorylated primer remained. We suspect that this leftover uncapped disphosphorylated primer is the reason we still see some product resulting from priming by a full-length (uncleaved) RNA primer.

We hypothesized that the size of the snatched fragment (here this is 15 nt) was dictated at least in part by the physical distance between the cap-binding site of the CBD and the active site of the ENDO. In theory, it should not matter how long the capped primer is provided it meets the minimum structural requirements, which are reported to be in the range of 12-18 nts (20). Thus, we should see the same cleaved-capped primer-incorporated product. To test this hypothesis, we repeated our *in vitro* cap-snatching assay with a shorter capped primer that was 24 nts long (Primer D, Supplementary Table 1) instead of 32 nts and detected an identical cleaved-capped primer-incorporated product for which the primer must have been cleaved to the same extent (Figure 1). To conclude, here, we demonstrate the full process of cap-dependent transcription initiation by a bunyavirus L protein *in vitro* starting with primer generation by cap-binding and ENDO activities, followed by cap-primed RNA synthesis by the RdRp.

### The SFTSV L protein transcription priming structure

Next, we sought to visualise by single-particle electron cryo-microscopy (cryo-EM) the conformational and structural changes associated with viral genome transcription by cap-snatching for the SFTSV L protein. To first visualise the SFTSV L protein in a transcription-specific priming state, we incubated the L protein with 5’ and 3’ RNA and a capped RNA primer (17 nts including the 5’ cap^0^ structure), which was in 6-fold molar excess to the 5’ and 3’ RNA species. To ensure that the L protein could not proceed to elongation, this reaction was set up in the absence of NTPs. This ultimately yielded a 3.35 Å resolution structure that we refer to as TRANSCRIPTION-PRIMING. Here, the 5’ and 3’ RNAs form the expected distal-duplex with the 3’ RNA feeding into the L protein core (Figure 2A). However, the domains at the N and C termini of the L protein have rearranged from their previously-determined positions (Figure 2A). Whereas, in our published replication-mode LATE-ELONGATION structure (19) the CBD is found in a peripheral location proximal to the ENDO, here the CBD is found at the base of the L protein. Bound to the CBD we found capped RNA, which as a result of the rearrangement of the ENDO and CBD is now oriented such that it feeds directly into the L protein core: 4 nts of the capped RNA 3’ end base-pair with the 3’ RNA template in the RdRp active site (Figure 2A; Supplementary Figure 3).

**Figure 2.**
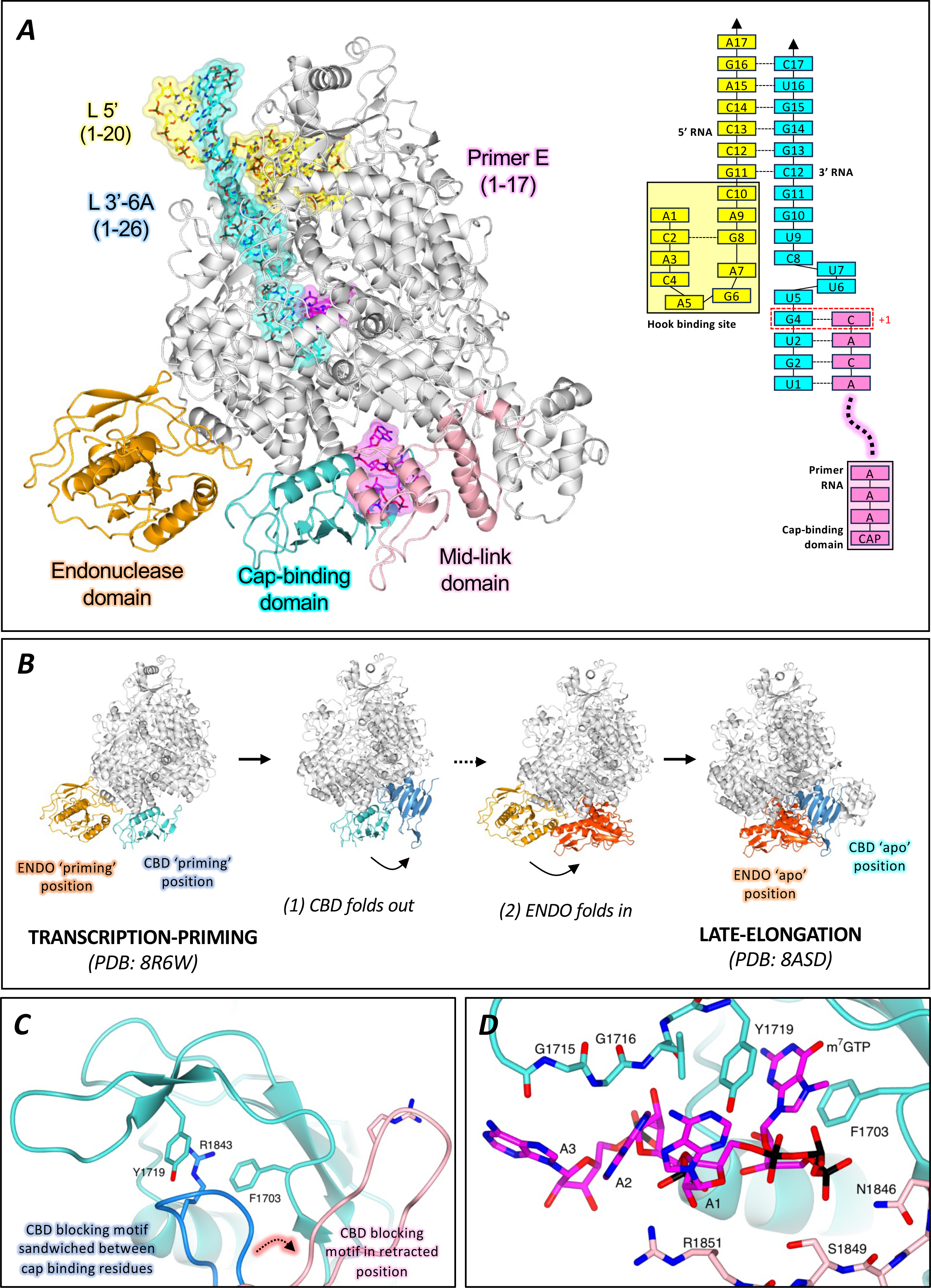
The SFTSV L protein transcription priming structure. (A) The SFTSV L protein (TRANSCRIPTION-PRIMING) is shown with key L protein domains coloured according to the following colour scheme: endonuclease (orange), cap-binding domain (cyan), and mid-link domain (light pink). RNAs are shown as sticks with surface overlaid (30% transparency) and coloured either yellow (5’ RNA), cyan (3’ RNA), or magenta (primer RNA). In addition, the RNAs present in each structure are shown schematically and labelled. (B) The conformational changes undergone by the L protein as it moves from transcription-priming to late-elongation are shown. L protein domains are coloured as in A. For clarity, RNA is not shown. (C) The position of the cap-binding domain (shown in cyan) blocking motif (composed of the Q1840 and R1843 sidechains) is shown sandwiched between the putative cap-binding residues (F1703 and Y1719) in the published LATE-ELONGATION structure (coloured dark blue) and in a retracted position in the TRANSCRIPTION-PRIMING structure published here (coloured light pink). (D) The cap-binding domain is shown with bound capped RNA. The cap-binding domain is coloured cyan, whereas the neighbouring mid-link domain is shown in light pink. The bound capped RNA is shown as sticks. Sidechains of key interacting amino acids, coloured according to the protein domain, are shown as sticks and labelled accordingly.

The ENDO domain has rotated by around 170° away from the base of the RdRp core in respect to its position in the *apo* conformation of L (18) (Figure 2B). Importantly, the movement of the ENDO domain creates sufficient space into which the CBD can then rotate by approximately 90° (Figure 2B). It is noteworthy that this position of the ENDO in the TRANSCRIPTION-PRIMING structure is significantly different to the ‘*raised conformation’* we reported in our genome replication-related EARLY-ELONGATION-ENDO structure (19). In total, we have now visualized the ENDO domain in three different conformations (Supplementary Figure 4). Interestingly, the endonuclease linker, which connects the endonuclease domain to the RdRp core does not appear to change significantly either its position or secondary structure (Supplementary Figure 4). The key appears to be the hinge region (residues 206 – 218) which connects the ENDO domain to the linker. This hinge region is largely unstructured in our previously published SFTSV L protein structures, with the exception of the EARLY-ELONGATION-ENDO structure (19). It appears therefore that this short stretch of amino acids imparts significant flexibility to the SFTSV L proteins endonuclease domain.

The outcome of these multi-domain conformational changes is that the putative cap-binding site is no longer oriented towards the solvent, as observed in our published LATE-ELONGATION structure (19), but instead oriented towards the RdRp core. In this conformation, bound capped RNA can proceed directly into the RdRp core to base-pair with 3’ RNA. Furthermore, the arginine finger blocking motif (composed of residues I1839 – V1844), that we and others found occluding access to the cap-binding site (19), has retracted with the blocking R1843 sidechain now sitting orthogonally to the cap-binding pocket (Figure 2C). Inspection of the map around the putative cap-binding site shows capped RNA bound with the guanosine moiety sandwiched between two aromatic sidechains: F1703 and Y1719 (Supplementary Figure 5). This mode of binding agrees with that described in the X-ray crystal structure of the SFTSV CBD in complex with the cap-analog m^7^GTP (18).

There are further stabilizing interactions between the cap structure and nearby residues. These include Q1707, which interacts with the cap guanosine, as well as K1668 and L1772, which coordinate the cap ribose via the O3’ and O2’ hydroxyls, respectively. There are extensive interactions between the cap^0^ triphosphate linker and residues from the C terminus of the L protein, including the sidechains of Y1719, N1834, N1846, and S1849. These sidechains interact with one or more phosphate oxygens and in doing so support cap-binding (Figure 2D). It is worth noting that the interactions with the triphosphate moiety seen here differ slightly from the published crystal structure, however, that is perhaps to be expected given that we are looking at the full L protein in the structures published here (18). Although the cap guanosine is bound perpendicular to the L protein core, the cap triphosphate linker points towards the RdRp active site. This is principally a result of two stretches of amino acids from the L protein C terminus, inclusive of M1845 – P1852 and T1838 – D1831, which direct the triphosphate linker and therefore the remaining nts attached to the cap structure towards the RdRp active site (Figure 2D). The quality of the map allows to confidently fit the first 3 nts of the capped RNA, and this corresponds to 5’-cap^0^-AAA-3’. A1 is supported largely by long interactions between the R1851 sidechain head group, which interacts with the A1 ribose, and the Y1719 sidechain which coordinates the A1 phosphate. There are interactions between A2 and surrounding amino acid residues. This includes the backbone nitrogen of V1717 and R1712 sidechain, which interact with the A2 ribose and base. There are effectively no protein-RNA interactions stabilizing the A3 base, although the A3 phosphate is shown to interact with the D1771 – G1773 backbone and it seems there are long interactions between the R1712 sidechain and the A3 N6/N7 (Figure 2D). Thus, it appears from the TRANSCRIPTION-PRIMING structure that there are limited protein sidechain-capped RNA base interactions which could indicate that the SFTSV L protein has little sequence specificity for the RNA snatched. A map of these interactions is provided in Supplementary Figure 6.

Similar to our published EARLY-ELONGATION structure (19), in the TRANSCRIPTION-PRIMING structure, the 5’ RNA binds in a hook-like conformation in a positively-charged cleft delineated by residues from the vRNA binding lobe (vRBL), fingers domain, and PA-C like core lobe. The 5’ and 3’ RNA then form a duplex stabilized by interactions between nts in the distal ends of the 5’/3’ RNAs which positions the 3’ RNA template in the RdRp active site ready for initiation. We can fit an unbroken chain of 3’ RNA nts from the distal duplex to the RdRp active site, allowing us to determine that the template is 4 nts into the active site with G4 at the +1 position (Supplementary Figure 3). Similar to that seen in our EARLY-ELONGATION structure (19), the 3’ RNA between the distal duplex and the RdRp active site is relatively condensed and appears to be anchored in place by the G10 base, which sits in a pocket delineated by W1342 – K1347 and L1399 – S1400 (Supplementary Figure 7). Further inspection of the map in the RdRp core itself led us to discover a 4 nts long RNA in the product position base-pairing with the 3’ RNA template (Supplementary Figure 3). Given that the L protein was incubated without any NTPs in this condition, it is not possible for this RNA to be an L protein synthesized product. We therefore propose these 4 nts correspond to the tail-end of the capped RNA, whose cap is bound to the CBD sitting at the base of the RdRp core.

The primer used in these experiments is designed to have a 4-bp overlap with the 3’ RNA template. In this schema, C1 of the primer base-pairs with G4 of the template RNA while A2 of the primer base-pairs with U3 of the template RNA (Supplementary Figure 3). This repeats with C3 and A4 of the primer which base-pair with G2 and U1 of the template RNA, respectively. The quality of the map in the TRANSCRIPTION-PRIMING structure is sufficient to fit the C1, A2, C3, and A4 of the primer with confidence (Supplementary Figure 3). C1 is supported by several long interactions which coordinate the C1 ribose O2’ (R920, Q1080, G1081) and O3’ hydroxyl groups (H1084). While A2 is coordinated by the S1125 sidechain, C3 is supported by interactions from the N1182 and R1197 sidechains. A4 is coordinated principally by the Q1204 and Q1224 sidechains. After A4, the map degrades to such an extent that fitting any further primer nts becomes speculative. It is noteworthy that while we can fit the 5’ and 3’ ends of the capped primer, we are missing effectively half of the primer nts from the map. This is likely because of the general lack of interacting sidechains to stabilize those nts as they travel from the CBD to the RdRp active site.

### The transition from cap-snatching to early-elongation

We next sought to capture the SFTSV L protein immediately after cap-snatching as it begins the process of primer elongation. For this, we incubated the L protein with nts 1-20 of the 5’ RNA, a modified 3’ RNA template (3’-midAAA) in addition to a 16 nt primer with a cap^0^ structure together with ATP, GTP, CTP, and a non-hydrolysable analog of UTP (NhUTP). The modification of the template, in which nts 6-8 are replaced with A’s, allowed us to stall the L protein *via* incorporation of NhUTP 2 nts downstream of the primer, which was designed to base-pair with nts 1-4 of the template RNA. This approach led to a single high-resolution structure, which we refer to as the TRANSCRIPTION-EARLY-ELONGATION (3.40 Å) structure (Figure 3A).

**Figure 3.**
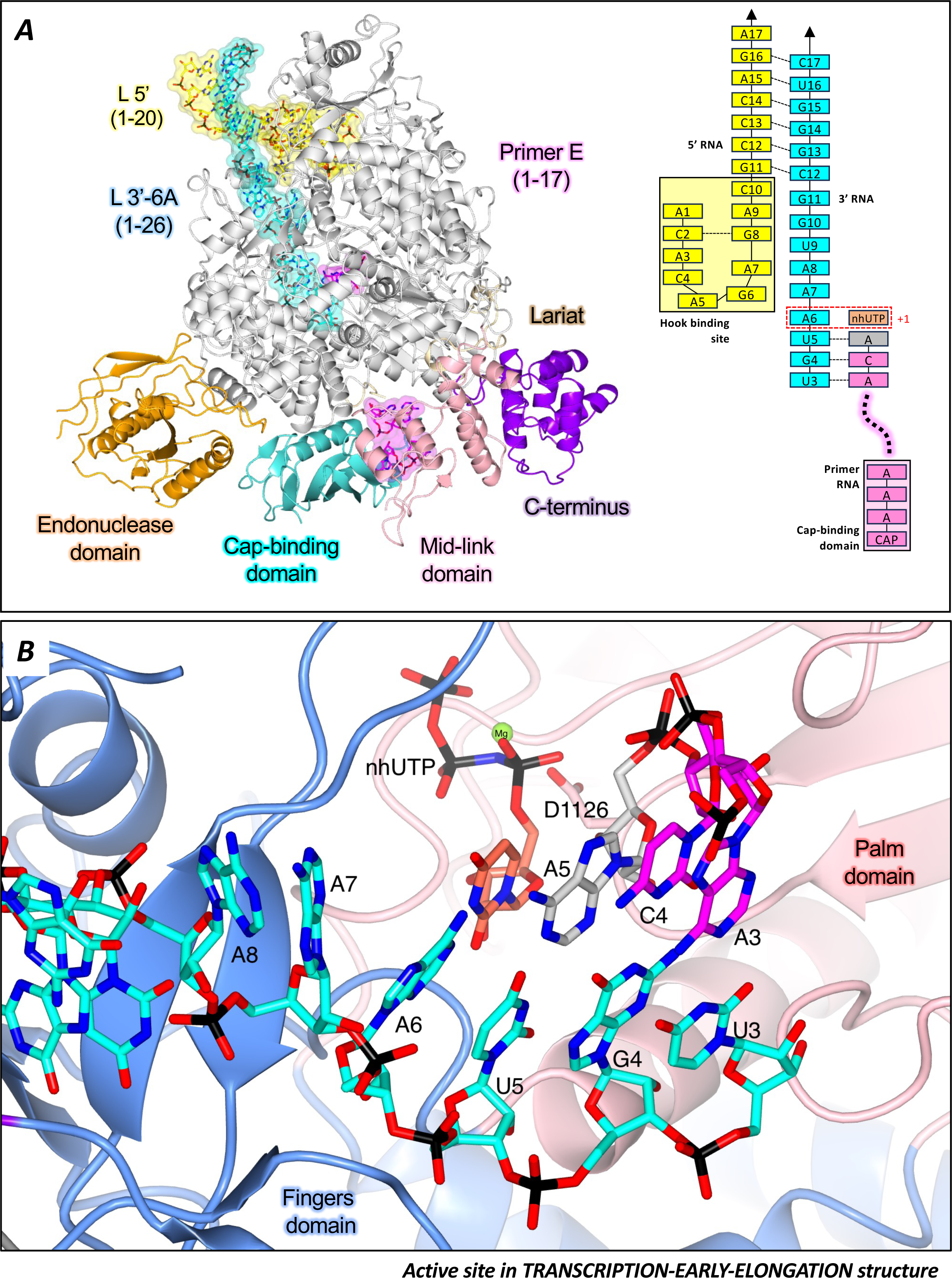
The transition from cap-snatching to early-elongation. (A) The SFTSV L (TRANSCRIPTION-EARLY-ELONGATION) is shown with key L protein domains coloured according to the following colour scheme: endonuclease (orange), cap-binding domain (cyan), and mid-link domain (light pink). RNAs are shown as sticks with surface overlaid (30% transparency) and coloured either yellow (5’ RNA), cyan (3’ RNA), or magenta (primer RNA). In addition, the RNAs present in each structure are shown schematically and labelled. (B) The active site in the TRANSCRIPTION-**EARLY-ELONGATION** structure is shown. Residues from the SFTSV L protein are labelled and coloured according to the assigned domain (blue for fingers domain, and coral for the palm domain). RNAs are shown as sticks and coloured as in A.

We note that the ENDO has moved by several angstroms away from the base of the L protein and the C terminus, which includes the so-called Lariat domain, has rotated towards the L protein core (Supplementary Figure 8). These movements correlate with the shunting of the capped primer (still bound by the CBD) by 2 nts as a result of the short elongation reaction while the 3’-5’ distal duplex region still stays intact. Furthermore, similar to the TRANSCRIPTION-PRIMING structure, we can fit an unbroken chain of 3’ RNA nts feeding directly from the distal duplex into the RdRp active site. However, herein lies a key difference. In the TRANSCRIPTION-PRIMING structure, at the splitting of the distal duplex, the 3’ RNA inverts at G10 and is then shown to ‘*bunch up’* immediately before entering the RdRp core. Instead, the 3’ RNA in the TRANSCRIPTION-EARLY-ELONGATION structure is shown to feed directly in to the RdRp core without inflection or bunching (Supplementary Figure 8). Further, despite the fact the map for these nts is poorly defined, it appears that these nts are under tension.

We also note that the inner L protein core has started to expand presumably to accommodate the growing product-template duplex, which we find to be only partly defined in the map. In the RdRp core, the quality of the map is sufficient to fit the terminal nucleotides 3-6 of the 3’ template RNA (5’ – A6-U5-G4-U3 – 3’) with A6 sitting at the +1 position (Figure 3B). Opposite A6, we can fit the stalling NhUTP, the tail of which is shown to support the coordination of a single Mg^2+^ ion alongside A986 and D1126. Adjacent to the NhUTP, we can fit 3 nts of the *‘product RNA’*, which in this structure is 5’-ACA-3’. The primer used in this reaction was designed to base-pair with nts 1-4 of the 3’ RNA. We conclude that the 5’-terminal A and C in this short product correspond to the last 2 nts in the capped primer, with the second A representing the first nt added by the L protein after priming. The L protein had successfully primed RNA synthesis using the capped-RNA provided and then proceeded to elongation at position 5 of the 3’ RNA. Stalling occurred at position 6 of the 3’ RNA template, which corresponds to the first A in this specifically designed template sequence. In comparison to the TRANSCRIPTION-PRIMING structure, the 3’ RNA template has been pulled forward by two nts and, as a result, the preceding nts feeding up to the distal duplex are now under tension. This tension, however, has not yet cause the distal duplex to dissolve.

It is noteworthy that we do not resolve the first two 3’ RNA nts or the two nts from the capped-RNA, to which they are base-paired in the map. We suggest this is principally due to the openness of the inner L protein core at this stage in the elongation process. There are very limited potential protein sidechains that could stabilize a growing product-template duplex and we clearly do not have the same number of RNA-RNA interactions that likely contribute to the stabilization of the 10-bp product-template duplex observed in our previously published replication-focussed LATE-ELONGATION structure (19). The growing product-template strands head towards the base of the L protein core, which remains buttressed by the CBD bound to the capped RNA. This is reasonably clear in our final polished cryo-EM map, where at lower contours the density for the product RNA connects to the density feeding from the bound cap-RNA fragment. We are confident therefore, that what we are seeing in this structure is the capped RNA bound simultaneously to the CBD, which remains in the priming position, and the 3’ RNA in the RdRp active site. We have chosen, however, not to fit the complete primer into the EM map, principally because of the poor nature of the map outside of the immediate primer termini. It is clear that the middle part of the primer retains significant flexibility adopting a range of orientations and we therefore cannot fit it with confidence.

In the TRANSCRIPTION-PRIMING and TRANSCRIPTION-EARLY-ELONGATION structures, we can now fit a significant portion of the extreme C terminus of the L protein, which thus far has been missing or poorly defined in other reported structural data. As a result, we can confidently fit the overwhelming majority of the L protein residues with only 4.3% of the entire L protein (90 amino acid residues) missing. There are further conformational changes to the L protein as we move from priming by cap-snatching to early-elongation of transcription, the most apparent being with respect to the CBD and Lariat domain (Supplementary Figure 9). Firstly, progression of the elongation reaction appears to have a shunting effect on the ENDO and CBD, which, when compared to the TRANSCRIPTION-PRIMING structure, are shown to recede slightly from itheir priming positions. Secondly, the C-terminal Lariat domain is shown to rotate and move towards the L protein core, presumably in response to the movement of the CBD. These domain movements are relatively fine and we think the reason for this can be explained by examining the inner L protein core. The internal L protein cavity is a relatively open space at this point in the elongation process. It is therefore entirely conceivable that the procession of the product-template strands by 2 nts (by comparison to the TRANSCRIPTION-PRIMING structure) can be largely ‘*absorbed’* by the openness of the L protein core. That being said, there is clearly a finite amount of space available. We expect the precession of the product-template strands to eventually be such that the CBD will need to move in a more significant manner, and ultimately returning to its ‘*apo’* position (19).

### Mutational analysis of capped RNA-interacting residues

The TRANSCRIPTION-PRIMING and TRANSCRIPTION-EARLY-ELONGATION structures have provided us with the first opportunity to identify amino acid residues involved in binding capped RNA in the cell. The capped RNA in the TRANSCRIPTION-PRIMING and TRANSCRIPTION-EARLY-ELONGATION structures appear coordinated by a mixture of sidechain-RNA phosphate and sidechain-RNA base interactions. The key residues of the CBD interacting with the m^7^GTP have already been tested previously (24). However, sidechains interacting with nts downstream of the m^7^GTP cap have overall not been tested. To investigate the importance of the additionally interacting sidechains, we mutated 19 different amino acids of the L protein and tested for genome replication and transcription activity in a cell-based viral mini-replicon system.

In a similar approach to that taken in Williams and Thorkelsson et al. (2023), we mapped the identified SFTSV residues onto the closely-related RVFV L protein to test the corresponding positions in our established RVFV mini-replicon system (Figure 4). Of the 19 mutants tested, 8 showed wild type-like activity (*i.e.*, Ren-Luc activity, >50%), including: V965G (SFTSV: N959), D1497A (SFTSV: Q1504), S1719A (SFTSV: T1709), V1726G (SFTSV: V1717), R1841A (SFTSV: R1829), S1861A (SFTSV: S1849), T1863A (SFTSV: R1851), and S1865A (SFTSV: R1853). Four mutants including, S1231A (SFTSV: N1224), E1565A (SFTSV: R1582), K1680A (SFTSV: K1668), and N1846A (SFTSV: N1834) demonstrated reduced L protein activity (*i.e.*, Ren-Luc activity, 8-49%) (Supplementary Table 3). We also found mutations to a small number of sidechains led to an inactive L protein phenotype (*i.e.*, Ren-Luc activity, 0-7%), including R1200A (SFTSV: R1193), W1205G (SFTSV: W1198), R1501A (SFTSV: R1499), T1563A (SFTSV: S1580), and H1858A (SFTSV: N1846) (Supplementary Table 3).

**Figure 4.**
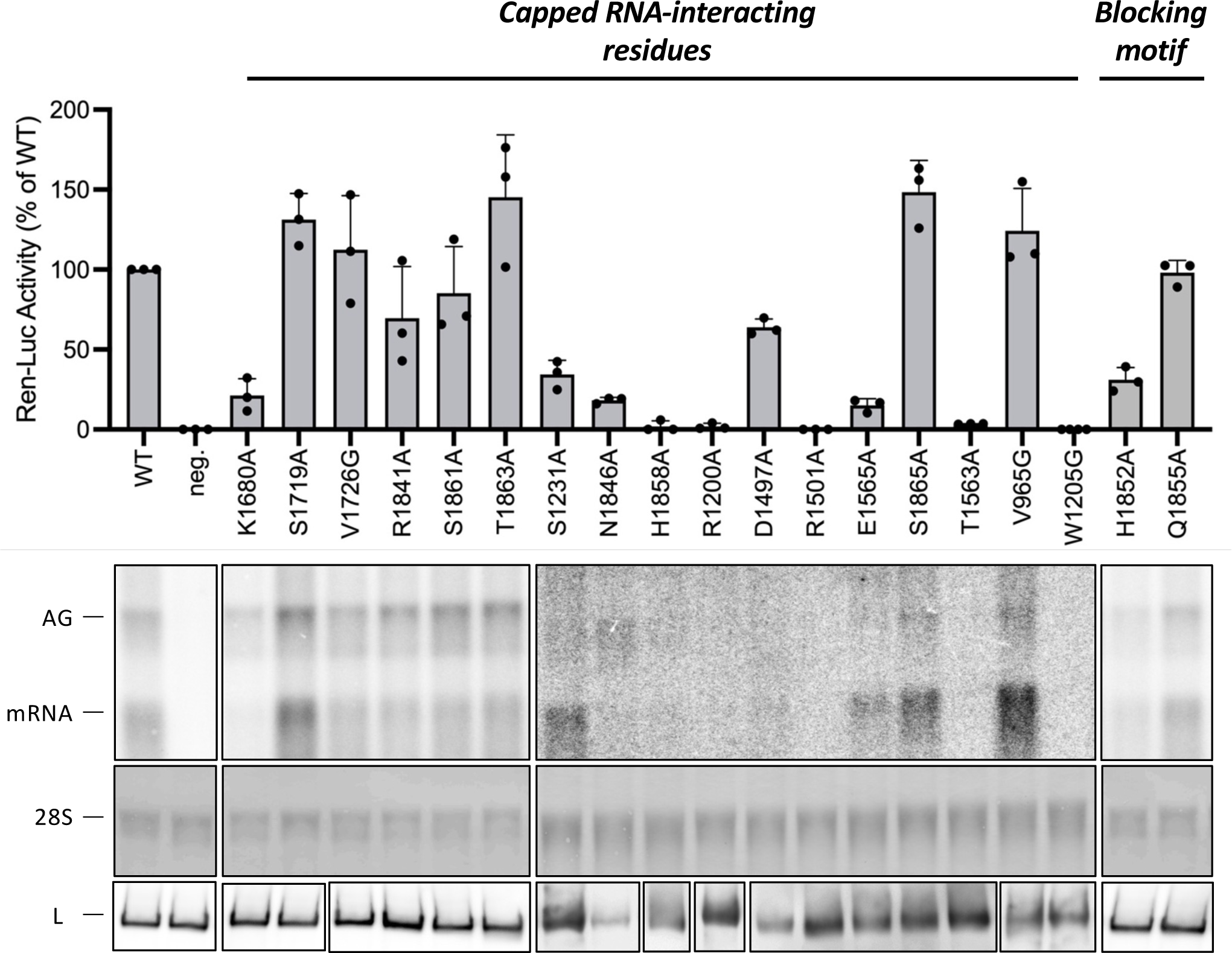
Mutational analysis of capped RNA-interacting residues. RVFV mini-replicon data for L protein with mutations to capped RNA-interacting residues presenting luciferase reporter activity (in standardized relative light units relative to the wild-type L protein (WT)). Data were presented as mean values ± SD of three biological replicates (n = 3). All biological replicates are shown as black dots (top panel). Middle panels present Northern blotting results with signals for antigenomic viral RNA (AG, equal to cRNA), viral mRNA (mRNA) and 28 S ribosomal RNA (28 S) as a loading control, and the bottom panel shows Western blot detection of FLAG-tagged L proteins (L) to demonstrate general expressibility of the mutants.

Of these inactivating mutations, the majority of the residues are located in the RdRp core and could possibly be involved in stabilizing the capped primer as it base-pairs with the 3’ RNA. As these residues line the walls of the RdRp core itself, it is also possible that altering the nature of the charged surface then impedes progression of the product-template duplex. Indeed, in previous studies, we have mutated adjacent residues thought to interact directly with the growing product-template duplex (*e.g.* SFTSV residues H1084 and F1085). and found that they also led to an inactive L protein (19). The only other mutant leading to an inactive L protein was H1858A (SFTSV: N1846), which coordinates the m^7^GTP α-phosphate. It is not entirely clear why mutating this residue has such a global effect but it is possible that it has a destabilizing effect.

Other mutations, like those to V1726, R1841, S1861, and T1863 are particularly interesting because it seems that the ∼50% reduction in the mRNA signal on the Northern blot is met with a 2-fold increase in the signal for antigenomic RNA (Supplementary Figure 9). This would suggest that disruption to these sites may inhibit conformational changes associated with transcription activity while leaving the L protein still capable of catalysing genome replication. This aligns with our structural insights into genome replication by the SFTSV L protein which does not appear to involve structural rearrangements to the C-terminal region (39). That being said, we could speculate a regulatory role for residues in the C-terminal domain in driving the balance between viral RNA genome replication and transcription. However, none of the mutants completely abolished the L proteins ability to undergo genome replication or transcription suggesting that the sidechains tested are not individually essential and do not critically impact viral transcription, genome replication or expressibility of the L protein.

We also know that under certain conditions the SFTSV cap-binding site is blocked by an arginine finger motif composed of the Q1840 and R1843 sidechains (Figure 2C). In our TRANSCRIPTION-PRIMING and TRANSCRIPTION-EARLY-ELONGATION structures, this motif is displaced by our capped primer but we nevertheless wanted to evaluate the importance of these two sidechains in our RVFV mini-replicon system (Figure 4). While the Q1855A (SFTSV: R1843) mutant showed a close to wild type-like phenotype, the H1852A (SFTSV: Q1840) mutant showed a largely reduced L protein activity (Supplementary Figure 9) (Supplementary Table 3). There was no selective effect on viral transcription detectable in the Northern blot. Thus, we concluded that the absence of this blocking motif may not directly lead to enhanced transcriptional activity. It is plausible, that the interaction of this blocker motif with the cap-binding site may primarily be a way for the L protein to stabilize a relatively flexible surface loop (including the blocker motif) rather than autoinhibiting the CBDs cap-binding capabilities.

### Structural insights into cap-independent primer elongation observed *in vitro*

In cells, we know that bunyaviruses, including SFTSV, must cap-snatch in order to produce their viral proteins. Translation is normally a cap-dependent process. It is curious therefore that, *in vitro*, we have also seen uncapped priming of RNA synthesis by SFTSV L protein (18,19). This is a phenomenon reported multiple times for different bunyaviruses. Indeed, for some bunyaviral L proteins it has been difficult to demonstrate cap-dependent primer elongation at all, such as LASV (34). However, it has not been clear how these different bunyaviral L proteins incorporate uncapped RNA primers into their growing strands. The **TRANSCRIPTION-PRIMING (*in vitro*)** structure (2.98 Å) we include here, provides the first structural insights into how this might happen in practice (Figure 5).

**Figure 5.**
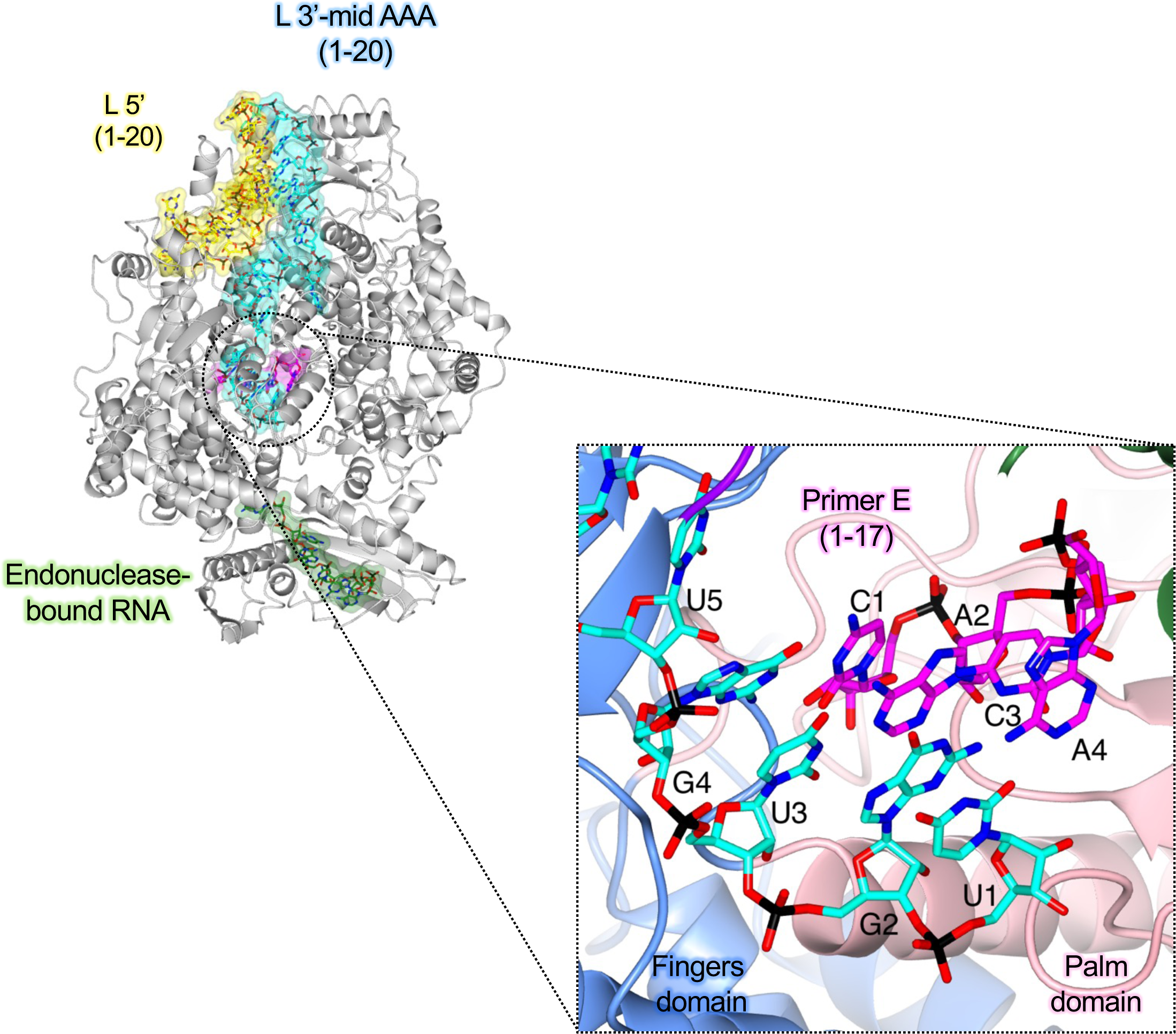
Structural insights into cap-independent primer elongation observed *in vitro*. (A) The SFTSV L (TRANSCRIPTION-EARLY-ELONGATION (*in vitro*)) is shown with bound RNAs shown as sticks with surface overlaid (30% transparency) and coloured either yellow (5’ RNA), cyan (3’ RNA), magenta (primer RNA), and sea green (endonuclease-bound RNA). A zoomed-in view of the active site in the TRANSCRIPTION-EARLY-ELONGATION (*in vitro*) structure is shown. Residues from the SFTSV L protein are labelled and coloured according to the assigned domain (blue for fingers domain, and coral for the palm domain). RNAs are shown as sticks and coloured as in A.

Here, the L protein adopts an overall similar conformation to that seen in our previously published EARLY-ELONGATION structure (19), a snapshot of the genome replication process. In the TRANSCRIPTION-PRIMING (*in vitro*) structure, the ENDO has returned to its ‘*apo’* position (19), now sitting across the base of the L protein core. For this structure, we specifically wanted to visualise transcription priming without allowing elongation to begin and so this reaction was incubated in the absence of NTPs. The 5’ RNA binds in a hook-like motif between the vRNA binding lobe (vRBL), fingers domain, and the PA-C like core lobe. The distal ends of the 5’ and 3’ RNAs form a distal-duplex and the quality of the map allows us to fit the 3’ RNA nts feeding from the distal duplex to the RdRp active site. We could determine therefore that, in a similar fashion to that seen in our EARLY-ELONGATION and EARLY-ELONGATION-ENDO structures (19), the 3’ RNA sits 4 nts into the L protein active site with G4 at the +1 position. There is density present for nts that would base-pair with the 3’ RNA in the L protein core. Given that no NTPs were provided for this reaction and that there is no incoming nucleotide at the +1 position, we propose this corresponds to nts from the RNA primer that have base-paired with the template RNA. We propose the TRANSCRIPTION-PRIMING (*in vitro*) structure corresponds to something fairly routinely *in vitro*, that is cap-independent primer elongation. This would explain why the CBD and ENDO domains are in their ‘*apo’* positions. The capping reaction with which we add a cap^0^ structure is not 100% efficient and therefore uncapped triphosphorylated primer is expected to remain in the reaction. We also know from our TRANSCRIPTION-EARLY-ELONGATION structure that the CBD and ENDO conformations associated with transcription priming are retained in the earliest stages of primer elongation. Finally, the lack of NTPs in this reaction would not even allow elongation to occur. We think it likely then that the primer/product RNA base-pairing with the 3’ RNA in the TRANSCRIPTION-PRIMING (*in vitro*) structure is in fact carryover uncapped triphosphorylated primer RNA. This would explain why the CBD does not appear to be involved in the same way that we see in the TRANSCRIPTION-PRIMING structure. Thus, what we assume to have here, is structural evidence for cap-independent priming, which we also observe in our *in vitro* assays.

In our TRANSCRIPTION-PRIMING structure, we demonstrate that capped RNA fragments are incorporated into RNA synthesis products as a result of significant domain reorganization, principally relating to the CBD and ENDO domains. However, that does not appear to be the case here in the TRANSCRIPTION-PRIMING (*in vitro*) structure. Instead, the reorganization of the L protein is far more nuanced and reminiscent of our EARLY-ELONGATION structure related to genome replication (19). Specifically, there is a rotation of the ENDO domain towards the L protein core, which provides space for part of the thumb ring domain (∼L1509 – K1577) to rotate away from the L protein core, moving into the space vacated by the ENDO domain (Supplementary Figure 10). These movements lead to an opening up of the putative product exit channel and we propose that it is *via* this channel that the non-capped primer RNA enters the L protein core, the terminal nts of which ultimately then base-pair with the 3’ template RNA (Supplementary Figure 10).

## DISCUSSION

Bunyaviruses need to snatch capped RNA primers for transcription priming in order to access the cellular translation machinery. This cap-snatching process is handled in large part by the viral L protein. Understanding how the L protein enables cap-snatching and therefore viral transcription is important not only to increase our fundamental understanding of bunyaviruses but also for the rational design of therapeutic interventions. Here, we establish an *in vitro* cap-snatching assay and report several high-resolution cryo-EM structures of the SFTSV L protein in the early-stages of viral transcription. Comparison of our TRANSCRIPTION-PRIMING structure to our previously published SFTSV L protein structures mimicking viral genome replication is particularly insightful (18,19). It demonstrates that, after binding to capped RNA, the N-terminal ENDO and C-terminal CBD of the SFTSV L protein undergo a significant conformational change (Fig. 2). In the genome replication-associated LATE-ELONGATION structure, the CBD is oriented such that bound capped RNA would be directed away from the RdRp (19). We suspect that the positioning of the CBD in this LATE-ELONGATION structure likely represents a fishing conformation by which the L protein can capture nearby host cellular mRNAs. The major domain reorganization in our TRANSCRIPTION-PRIMING structure ultimately repositions the capped RNA-bound CBD such that it then sits across the floor of the L protein (Figure 6). This enables the CBD-bound capped RNA to feed directly into the RdRp core and base-pair with the 3’ RNA template. Addition of NTPs spiked with a stalling NhUTP that stalls 2 nts downstream of the incorporated primer shows that this transcription priming state is retained in our TRANSCRIPTION-EARLY-ELONGATION structure suggesting that perhaps it is the progression of the growing product-template duplex in the RdRp core that is ultimately responsible for shunting the ENDO and CBD back into their late-stage elongation positions (Figure 2B and 6).

**Figure 6.**
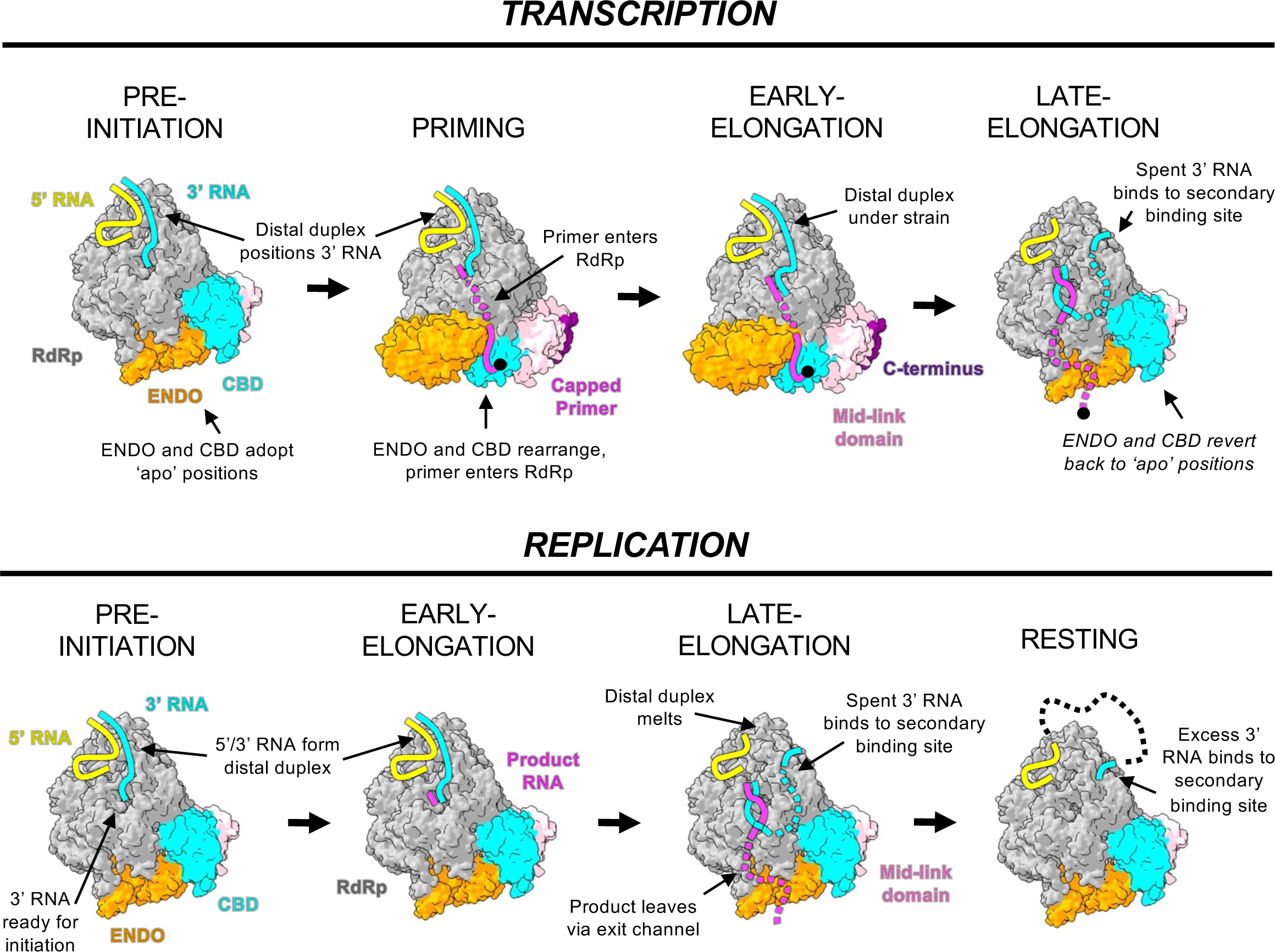
Structure-based models of SFTSV replication and transcription. L protein structures are displayed as surface with key domains coloured according to the following colour scheme: endonuclease (orange), cap-binding domain (cyan), mid-link domain (light pink), and C-terminus (purple). The remaining protein is coloured grey. RNAs are represented by coloured solid/dashed lines and coloured either yellow (5’ RNA), cyan (3’ RNA), magenta (product/primer RNA). Important to note that we consider the shift between early- and late-elongation to be characterised by the splitting of the product-template duplex in the RdRp core and subsequent exit of these RNA species via the product and template exit channels, respectively. Key features are indicated.

The determination of several transcription-specific state structures for the SFTSV L proteins allows to make comparisons to LACV L protein, the only other L protein for which transcription-state structures are available so far (33). In LACV, transcription initiation depends on a closure of the C-terminal region and major repositioning of the N-terminal ENDO. These conformational changes were suggested to prompt primer cleavage and primer entry into the RdRp core (39). While transcription initiation in SFTSV also includes a significant repositioning of the N-terminal ENDO, we observe that the C-terminal CBD is also highly mobile. There are therefore differences in how the L proteins from different bunyavirus families enable viral genome replication and transcription.

In our previously published LATE-ELONGATION state structure, although we could fit the CBD with confidence, the cap-binding site was blocked by an arginine blocking motif composed of residues T1838 – V1844 as also observed by others (51–53). In the TRANSCRIPTION-PRIMING and TRANSCRIPTION-EARLY-ELONGATION structures shown here, the blocking motif has been displaced by the capped RNA and now sits orthogonally to the CBD. Interestingly, whereas in the genome replication-related LATE-ELONGATION structure, this arginine motif presumably blocks access to the cap-binding site, in our TRANSCRIPTION-PRIMING and TRANSCRIPTION-EARLY-ELONGATION structures, the residues composing the blocking motif form a wall directing bound capped RNA towards the RdRp core. This is a nice example of multifunctionality, a quintessential feature of the bunyavirus multidomain L protein, which itself is responsible for enabling not just viral genome replication but also transcription.

The capped RNA is bound in principle as we expected with the m^7^GTP cap moiety sandwiched between two aromatic sidechains: F1703 and Y1719 (Fig. 2D). This follows nicely the mode of recognition seen in the crystal structure of the isolated SFTSV CBD bound to m^7^GTP cap-analog (18). What is of particular interest, however, is the coordination of nts downstream of the cap. The majority of sidechains supporting the coordination of the capped RNA fragment come from other domains than the CBD. Further, most sidechains interacted with the capped RNA *via* its phosphate backbone rather than with the nt base. This would indicate to us that capped RNA is bound by the L protein regardless of the exact sequence although we cannot exclude that the size of the base (purine vs. pyrimidine) may have an impact.

That being said, the capped RNA used in these experiments was modified by the addition of a cap0 structure. However, in cells, the most abundant cap structure is cap1, in which the first nt downstream of the m^7^GTP cap is additionally methylated (CH_3_) at the 2’-O position of the ribose (54,55). In our TRANSCRIPTION-PRIMING and TRANSCRIPTION-EARLY-ELONGATION structures, a 2’-O methyl group would be prohibitively close to the sidechain of R1851 and the base of the next RNA nt, which in our structures is an A (Supplementary Figure 11). This methyl/base clash would be less severe however, and perhaps not prohibitive at all, if the next base was smaller, *i.e.* a pyrimidine. Thus, for coordination of cap1-RNA, there may be a certain degree of sequence specificity or at least preference for a specific set of bases at position 2 after the m^7^GTP in the RNA sequence. Interestingly, a preference for smaller bases at this position was found in sequencing data of viral transcripts for the closely-related RVFV. In this case, it has been shown that the L protein preferentially snatches 5’-TOP mRNAs, which have a >75% chance of containing a pyrimidine at position 2 after the m^7^GTP (56). Of course, it is also possible that the R1851 sidechain rotates away from the bound capped RNA and in doing so creates space for the cap1 methyl group (Supplementary Figure 11). Flexibility in the RNA phosphate backbone may also provide a perfectly sensible solution to this problem. Further work is certainly needed to test these hypotheses and make any concrete proposals regarding the underlying molecular mechanism.

This does not mean however that the ENDO cannot impose some degree of sequence specificity downstream of the cap structure by virtue of preferentially binding and then cleaving specific RNA sequences or motifs. Indeed, it has previously been proposed based on biochemical studies that the phenuivirus endonuclease prefers purine-rich RNA species as substrates (25). Structural evidence lending weight to this argument is however lacking. Indeed, it would make sense for the ENDO to preferentially cleave at positions producing capped fragments ending in AC as this would then allow the capped fragment to base-pair with the SFTSV 3’ genome termini, which end with UG repeats. That being said, we do not see any strong evidence to suggest that the SFTSV L protein ENDO has a sequence specificity. There is a general lack of structural data showing the SFTSV ENDO bound to an RNA substrate. The only example of this can be found in our previously published LATE-ELONGATION structure, in which we show excess 5’ RNA bound across the ENDO active site. Even here, we could not identify any clear structural determinants that would indicate the ENDO would produce fragments ending in, for example, AC. Although we did speculate that perhaps residue Y76 may make the coordination of a bulky purine base in the equivalent of the P0 pocket (using influenza virus endonuclease nomenclature) less favorable.

Considering the important role the ENDO plays in the cap-snatching process, understanding how the ENDO is regulated is of particular interest. In a previously published structure, our EARLY-ELONGATION-ENDO structure, we demonstrated that the ‘*raised’* conformation adopted by the ENDO could be one example of such regulatory mechanism. In this conformation, access to the active site is effectively blocked by the positioning of the C-terminal α-helix of the SFTSV ENDO (residues 211 – 233) across the ENDO active site. Other bunyaviruses have been shown to adopt several different mechanisms for regulating their ENDO domain adding layers of redundancy and complexity to the ENDO regulation. For example, the LASV ENDO has been shown to be autoinhibited by two alternative mechanisms: either an inhibitory peptide or the C-terminal helix of the ENDO blocks the ENDO active site (34). In our TRANSCRIPTION-PRIMING and TRANSCRIPTION-EARLY-ELONGATION structures described here, we find that the C-terminal α-helix of the SFTSV ENDO blocking substrate access to the ENDO active site, albeit in a different way to that seen in the EARLY-ELONGATION-ENDO structure. We have therefore observed the SFTSV ENDO in three unique positions (sup fig. 3) and likely have identified two different mechanisms by which the access to the SFTSV ENDO active site is regulated.

Altogether, the biochemical and structural data provided here give a first glimpse into how the SFTSV L protein likely recognizes and binds to capped RNA fragments in infected cells. We show that the L protein reorganizes its N- and C-terminal domains significantly to present snatched fragments to the RdRp core. This demonstrates, quite unexpectedly, that this priming conformation is retained in the early stages of primer elongation meaning that the L protein adopts very different conformations at early elongation for viral genome replication and transcription. Future work will be needed to elucidate further steps of the viral genome replication and transcription mechanisms, such as termination. Nevertheless, by integrating the now quite significant number of high-resolution cryo-EM structures of the SFTSV L protein undergoing both genome replication and transcription with biochemical data and significant mutational analyses, we can build a more complete picture of how these processes are enabled at the molecular level (Figure 6).

## DATA AVAILABILITY

Coordinates and maps included in this paper have been deposited in the Worldwide Protein Data Bank (wwPDB) and the Electron Microscopy Data Bank (EMDB) with the following accession codes: SFTSV L protein in a transcription priming state with CBD bound to capped-RNA [TRANSCRIPTION-PRIMING] EMD-18967 PDB-8R6W; SFTSV L protein captured at early elongation following capped-primer incorporation [TRANSCRIPTION-EARLY-ELONGATION] EMD-18969 PDB-8R7Y; SFTSV L protein in a state reflecting cap-independent transcription priming [TRANSCRIPTION-PRIMING (*in vitro*)] EMD-18963 PDB-8R6U.

## ACKNOWLEDGEMENTS

We want to thank Stephan Günther for his support and helpful discussions throughout the project. We acknowledge support by Carolin Seuring, Cornelia Cazey and Ulrike Laugks for access to the CryoEM multi-user Facility at CSSB and providing time for sample preparation, screening, and data collection; Wolfgang Lugmayr for help and support in using the CSSB partition on the DESY computer cluster for cryo-EM data processing. We also want to thank Lennart Sänger for feedback on the manuscript. Finally, we thank the Wilhelm und Maria Kirmser-Stiftung for support.

## SUPPLEMENTARY TABLES AND TABLE LEGENDS

**Supplementary Table 1.**
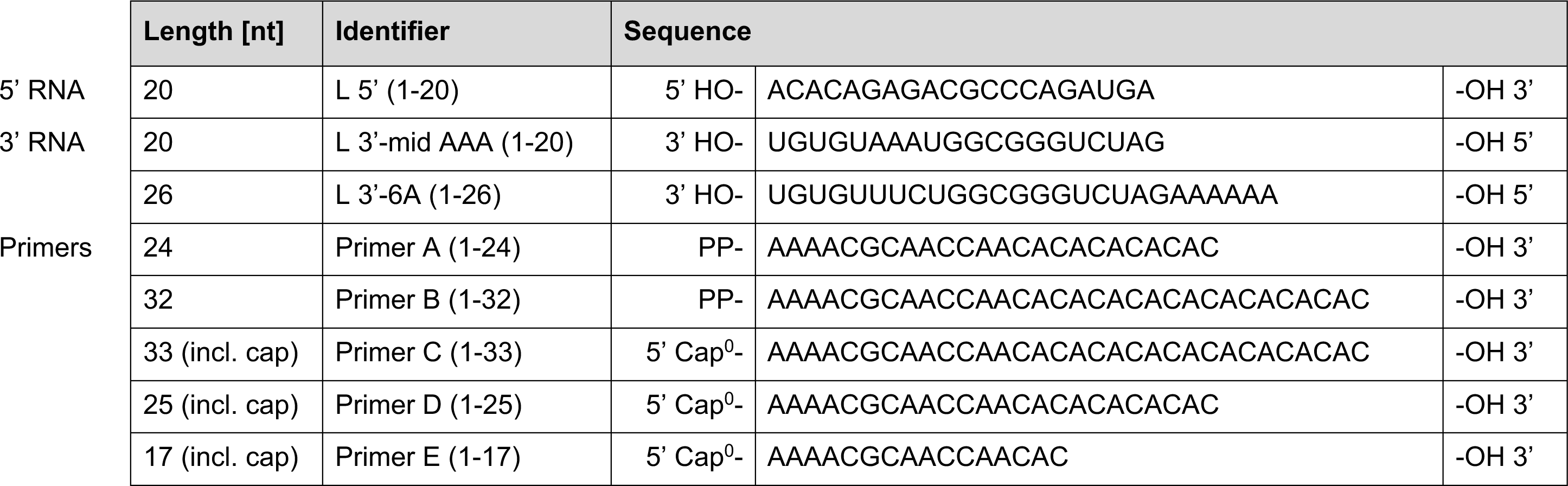
Synthetic RNA oligonucleotides used in different assays. This table lists all the RNA oligonucleotides used in this study. The sequence, length in nucleotides (nt) and the specific identifier used to label the RNA in the experimental descriptions are provided. All RNAs were chemically synthesized by Biomers with the exception of the primer RNA, which was chemically synthesized by ChemGenes and then manually capped using the ScriptCap kit (see Materials and Methods).

**Supplementary Table 2.**
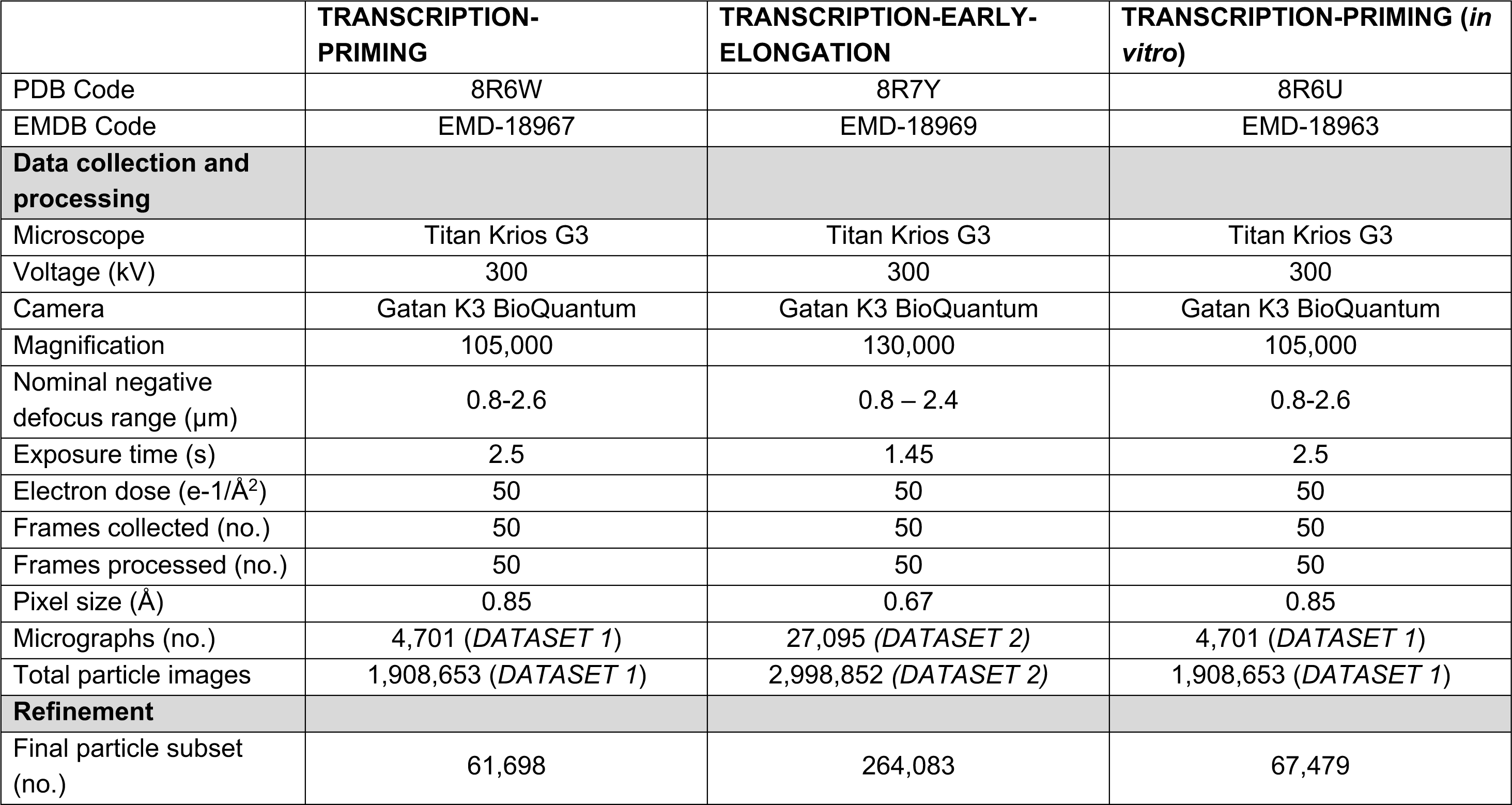
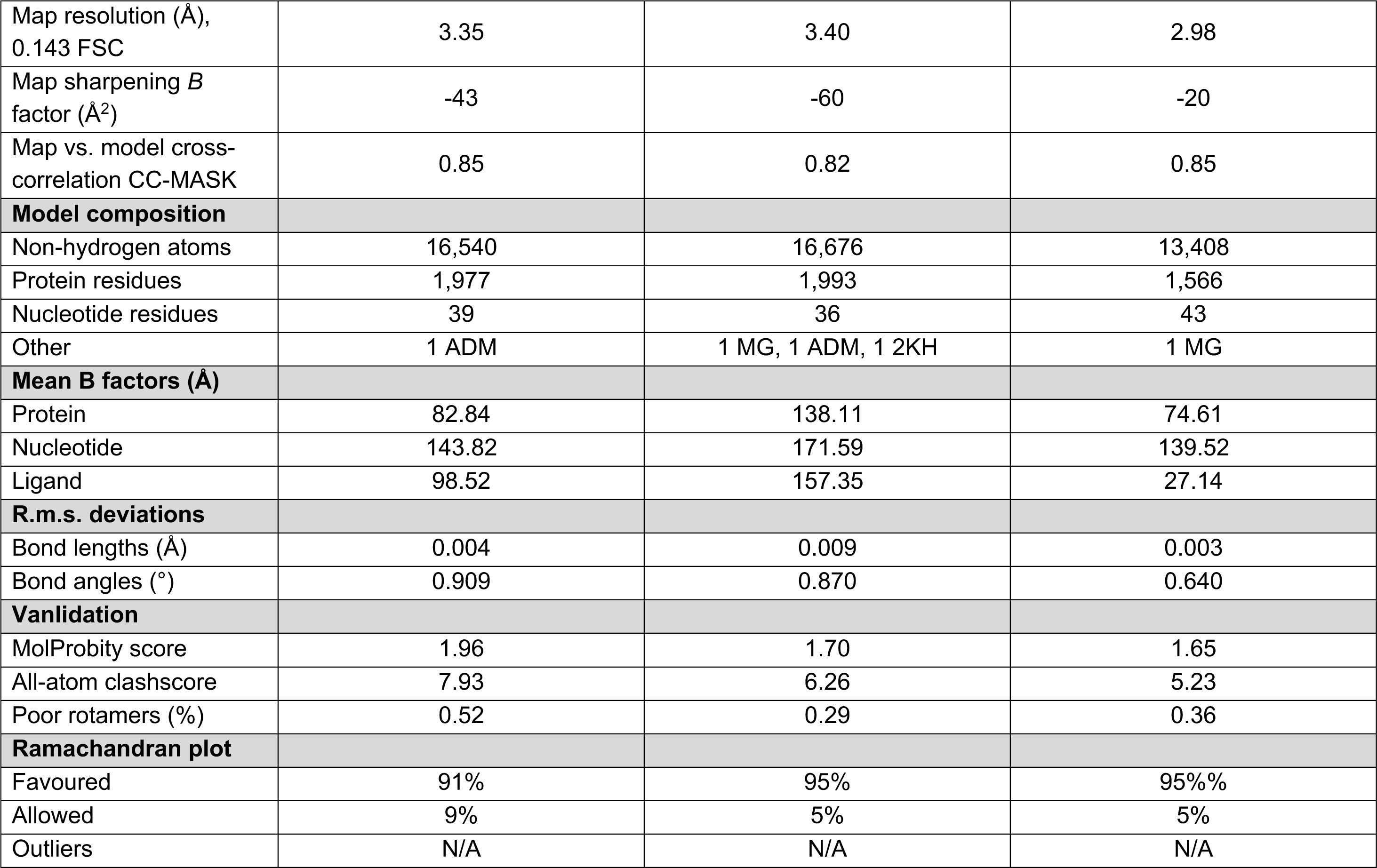
Cryo-EM data collection, processing, refinement, and validation statistics. This table provides the parameters and statistics for the data collection, processing, refinement, and structure validation of the cryo-EM structures described. Refinement statistics were generated using the Phenix package.

|  | **TRANSCRIPTION-PRIMING** | **TRANSCRIPTION-EARLY-ELONGATION** | **TRANSCRIPTION-PRIMING (*in vitro*)** |
| --- | --- | --- | --- |
| PDB Code | 8R6W | 8R7Y | 8R6U |
| EMDB Code | EMD-18967 | EMD-18969 | EMD-18963 |
| **Data collection and processing** |  |  |  |
| Microscope | Titan Krios G3 | Titan Krios G3 | Titan Krios G3 |
| Voltage (kV) | 300 | 300 | 300 |
| Camera | Gatan K3 BioQuantum | Gatan K3 BioQuantum | Gatan K3 BioQuantum |
| Magnification | 105,000 | 130,000 | 105,000 |
| Nominal negative defocus range ($\mu\text{m}$) | 0.8-2.6 | 0.8 – 2.4 | 0.8-2.6 |
| Exposure time (s) | 2.5 | 1.45 | 2.5 |
| Electron dose ($\text{e}^-/\text{\AA}^2$) | 50 | 50 | 50 |
| Frames collected (no.) | 50 | 50 | 50 |
| Frames processed (no.) | 50 | 50 | 50 |
| Pixel size ($\text{\AA}$) | 0.85 | 0.67 | 0.85 |
| Micrographs (no.) | 4,701 (*DATASET 1*) | 27,095 (*DATASET 2*) | 4,701 (*DATASET 1*) |
| Total particle images | 1,908,653 (*DATASET 1*) | 2,998,852 (*DATASET 2*) | 1,908,653 (*DATASET 1*) |
| **Refinement** |  |  |  |
| Final particle subset (no.) | 61,698 | 264,083 | 67,479 |
| Map resolution (Å),<br>0.143 FSC | 3.35 | 3.40 | 2.98 |
| Map sharpening *B*<br>factor (Å<sup>2</sup>) | -43 | -60 | -20 |
| Map vs. model cross-<br>correlation CC-MASK | 0.85 | 0.82 | 0.85 |
| **Model composition** |  |  |  |
| Non-hydrogen atoms | 16,540 | 16,676 | 13,408 |
| Protein residues | 1,977 | 1,993 | 1,566 |
| Nucleotide residues | 39 | 36 | 43 |
| Other | 1 ADM | 1 MG, 1 ADM, 1 2KH | 1 MG |
| **Mean B factors (Å)** |  |  |  |
| Protein | 82.84 | 138.11 | 74.61 |
| Nucleotide | 143.82 | 171.59 | 139.52 |
| Ligand | 98.52 | 157.35 | 27.14 |
| **R.m.s. deviations** |  |  |  |
| Bond lengths (Å) | 0.004 | 0.009 | 0.003 |
| Bond angles (°) | 0.909 | 0.870 | 0.640 |
| **Vanlilation** |  |  |  |
| MolProbity score | 1.96 | 1.70 | 1.65 |
| All-atom clashscore | 7.93 | 6.26 | 5.23 |
| Poor rotamers (%) | 0.52 | 0.29 | 0.36 |
| **Ramachandran plot** |  |  |  |
| Favoured | 91% | 95% | 95%% |
| Allowed | 9% | 5% | 5% |
| Outliers | N/A | N/A | N/A |

**Supplementary Table 3.** Raw data from RVFV mini replicon assays. For each mutant, the calculated Ren-Luc activity (as a % compared to the WT) from 3 biological replicates (as indicated in the table), the calculated average and standard deviation is provided.

| RVFV Residue | Replicate 1 | Replicate 2 | Replicate 3 | Average | Standard Deviation |
| --- | --- | --- | --- | --- | --- |
| **RNA-interacting residues** |  |  |  |  |  |
| K1680A | 11.65 | 32.32 | 20.02 | 21.33 | 10.40 |
| S1719A | 114.89 | 147.45 | 131.45 | 131.26 | 16.28 |
| V1726G | 111.49 | 78.92 | 146.70 | 112.37 | 33.90 |
| R1841A | 42.88 | 105.57 | 60.21 | 69.55 | 32.37 |
| S1861A | 70.93 | 65.76 | 118.90 | 85.20 | 29.30 |
| T1863A | 101.63 | 157.96 | 176.30 | 145.30 | 38.91 |
| S1231A | 24.98 | 35.71 | 42.49 | 34.39 | 8.83 |
| N1846A | 16.08 | 19.38 | 19.18 | 18.21 | 1.85 |
| H1858A | 0.03 | 0.03 | 5.92 | 1.99 | 3.40 |
| R1200A | 0.81 | 0.86 | 4.16 | 1.94 | 1.92 |
| D1497A | 59.93 | 69.73 | 62.07 | 63.91 | 5.15 |
| R1501A | 0.02 | 0.05 | 0.03 | 0.03 | 0.02 |
| E1565A | 16.74 | 10.44 | 18.11 | 15.10 | 4.09 |
| S1865A | 125.96 | 155.97 | 163.30 | 148.41 | 19.78 |
| T1563A | 3.56 | 3.28 | 3.58 | 3.47 | 0.17 |
| V656G | 109.92 | 154.89 | 107.90 | 124.24 | 21.69 |
| W1205G | 0.02 | 0.04 | 0.02 | 0.03 | 0.01 |
| **Blocking motif residues** |  |  |  |  |  |
| H1852A | 29.80 | 39.27 | 24.15 | 26.98 | 4.00 |
| Q1855A | 102.60 | 102.41 | 89.00 | 98.00 | 7.80 |

**Supplementary Figure 1.**
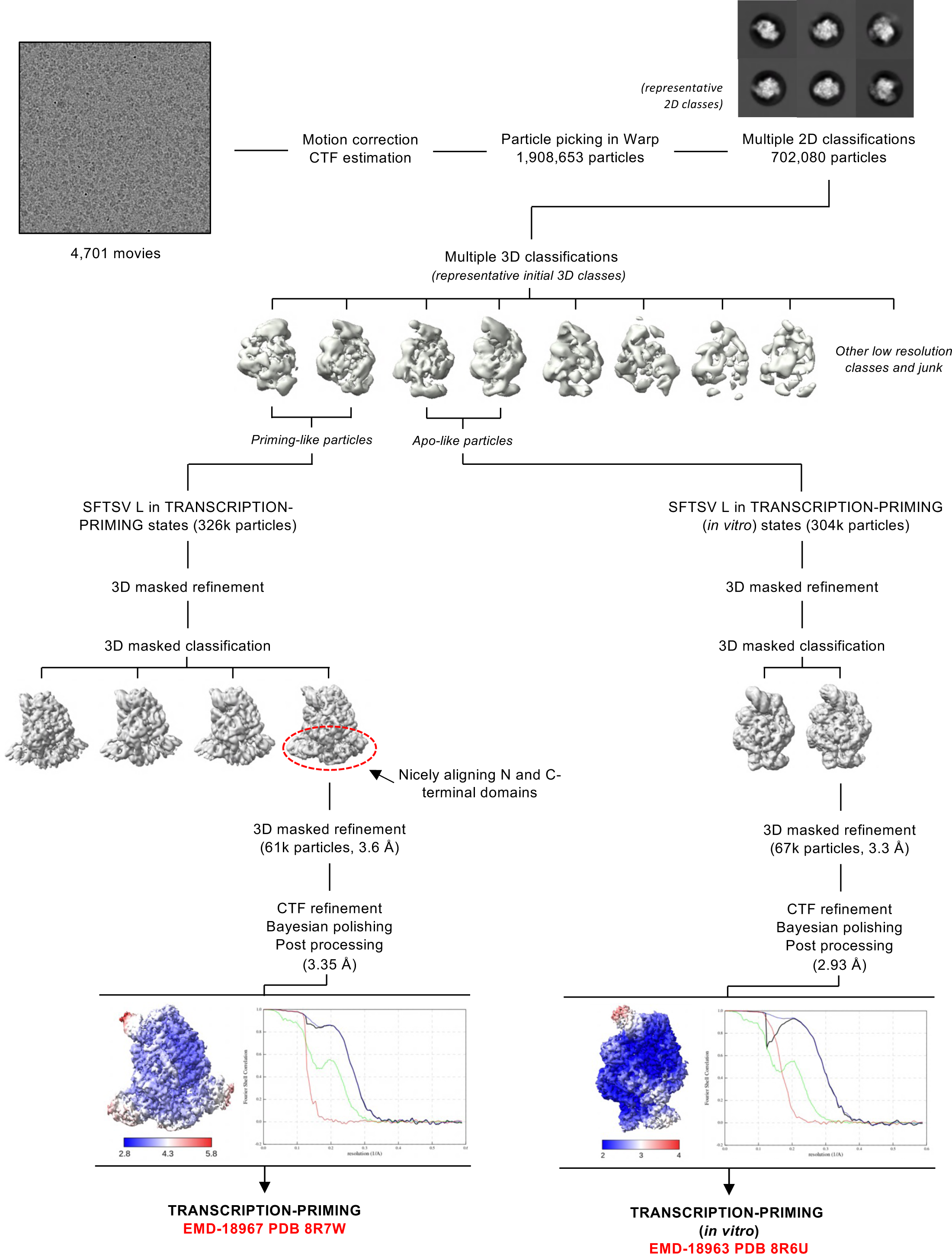
Image processing strategy to obtain TRANSCRIPTION-PRIMING and TRANSCRIPTION-PRIMING (in vitro) structures. Schematic representation of the image processing strategy used with the data collected on a Titan Krios G3 equipped with a Gatan K3 BioQuantum detector to obtain the **TRANSCRIPTION-PRIMING** and **TRANSCRIPTION-PRIMING (*in vitro*)** structures. Representative micrographs, 2D class averages, and 3D class averages are displayed. All processing steps, including 2D classification, 3D classification, 3D refinement, and post-processing were completed in RELION with the exception of the **TRANSCRIPTION-PRIMING** structure, the map for which was further processed in LocScale in CCP-EM. Local resolution EM maps are coloured according to the resolution key shown. Fourier shell correlation curves (FSC) curves are also displayed.

**Supplementary Figure 2.**
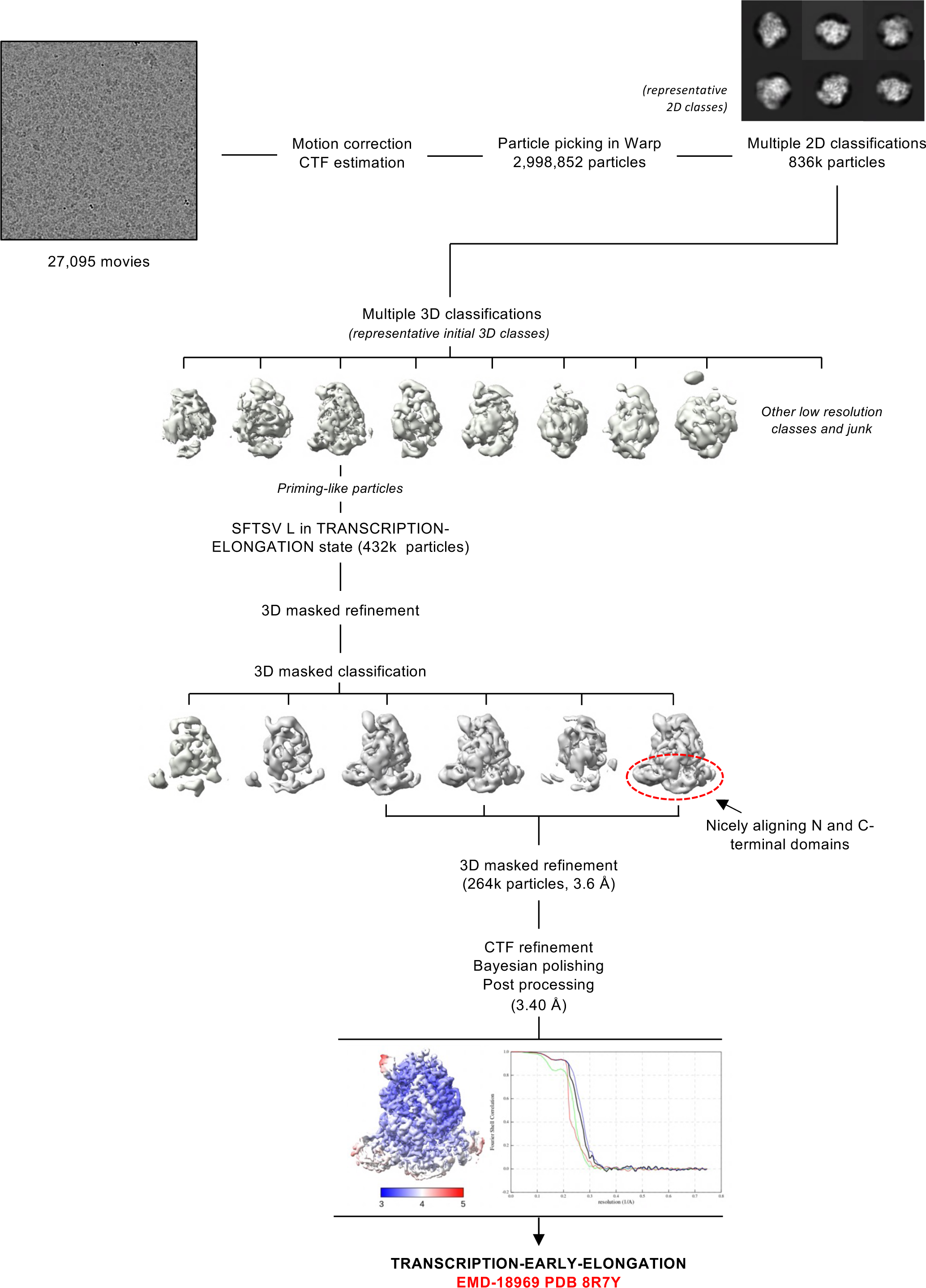
Image processing strategy to obtain TRANSCRIPTION-EARLY-ELONGATION structure. Schematic representation of the image processing strategy used with the data collected on a Titan Krios G3 equipped with a Gatan K3 BioQuantum detector to obtain the **TRANSCRIPTION-EARLY-ELONGATION** structure. Representative micrographs, 2D class averages, and 3D class averages are displayed. All processing steps, including 2D classification, 3D classification, 3D refinement, and post-processing were completed in RELION. Local resolution EM maps are coloured according to the resolution key shown. Fourier shell correlation curves (FSC) curves are also displayed.

**Supplementary Figure 3.**
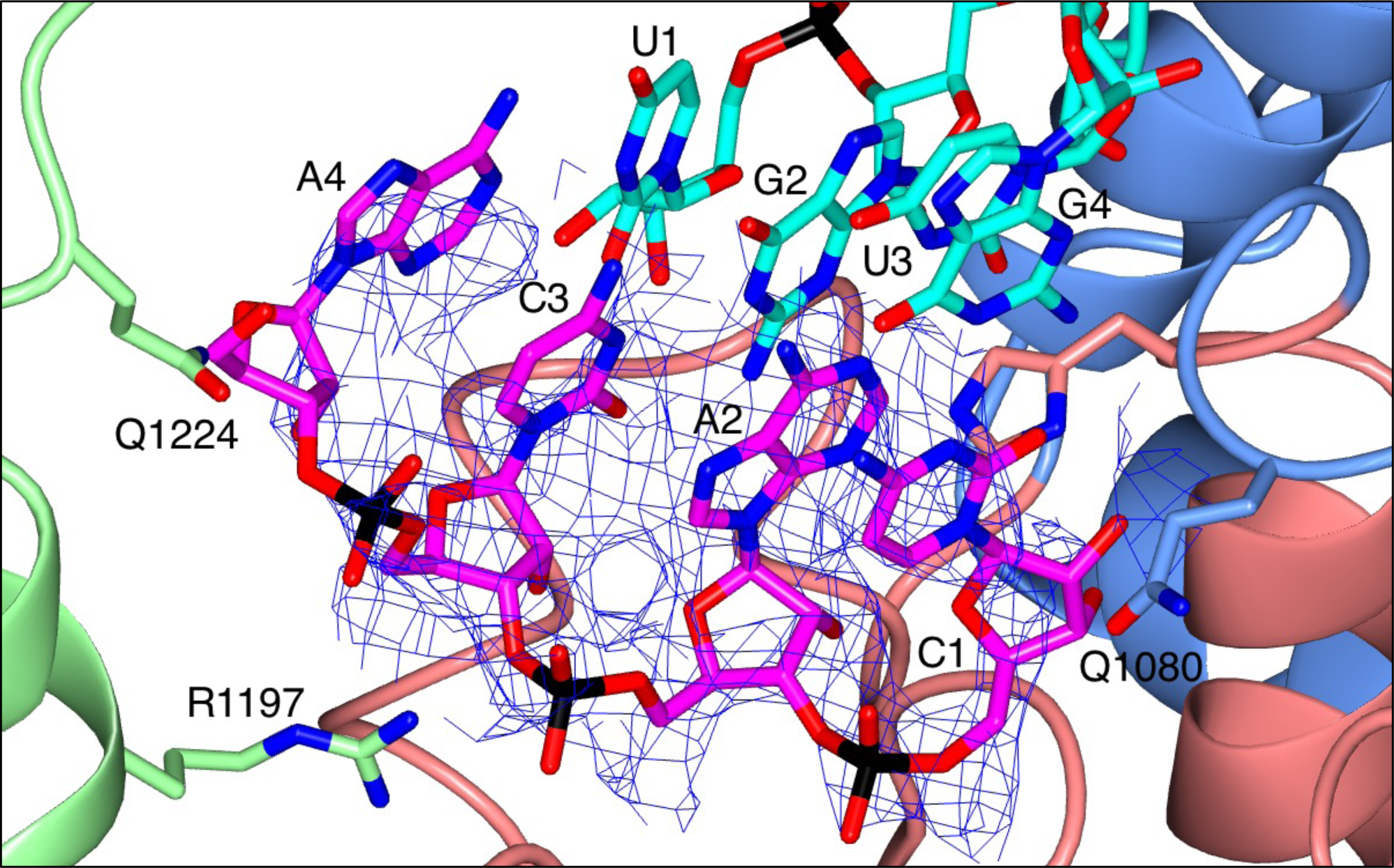
Zoomed-in view of the TRANSCRIPTION-PRIMING structure active site. A zoomed-in view of the active site in the **TRANSCRIPTION-EARLY-ELONGATION** structure is shown with the respective map overlaid. Residues from the SFTSV L protein are labelled and coloured according to the assigned domain (blue for fingers domain, light green for the thumb and thumb ring domains, and coral for the palm domain). RNAs are shown as sticks and coloured either yellow (5’ RNA) or magenta (primer RNA). The overlaid map is clipped to the primer RNA.

**Supplementary Figure 4.**
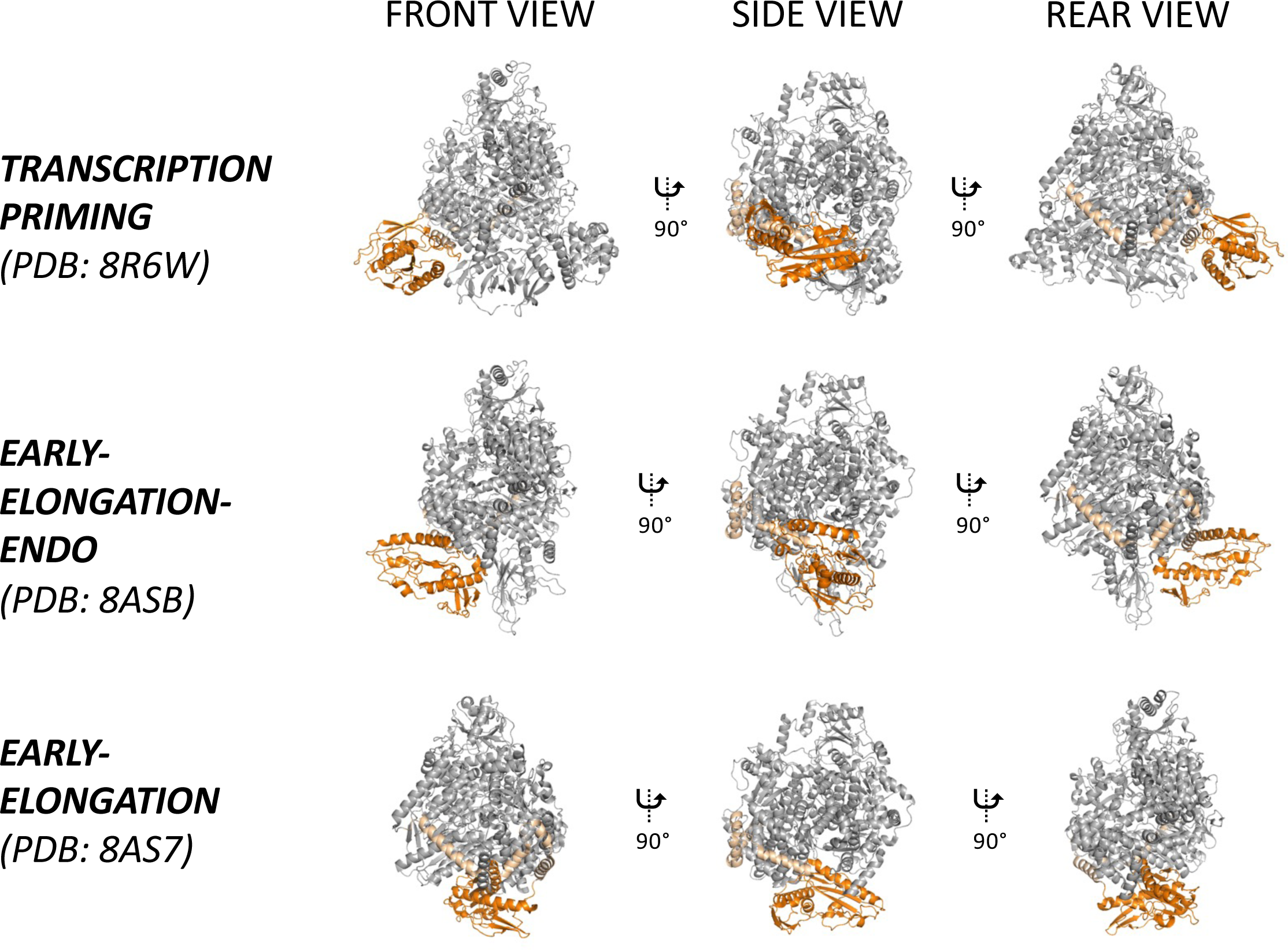
Hyper-flexibility of the SFTSV L protein endonuclease domain. The SFTSV L protein TRANSCRIPTION-PRIMING, EARLY-ELONGATION (PDB: 8AS7), and EARLY-ELONGATION-ENDO (PDB: 8ASB) structures are shown having been superposed by a least-squares fit of main-chain RNA-dependent RNA polymerase residues. For clarity, the majority of the L protein is coloured in light grey with only the endonuclease and endonuclease linker coloured orange and wheat, respectively. Bound RNA in each structure is not shown. Three views are shown for each structure to enable comparison.

**Supplementary Figure 5.**
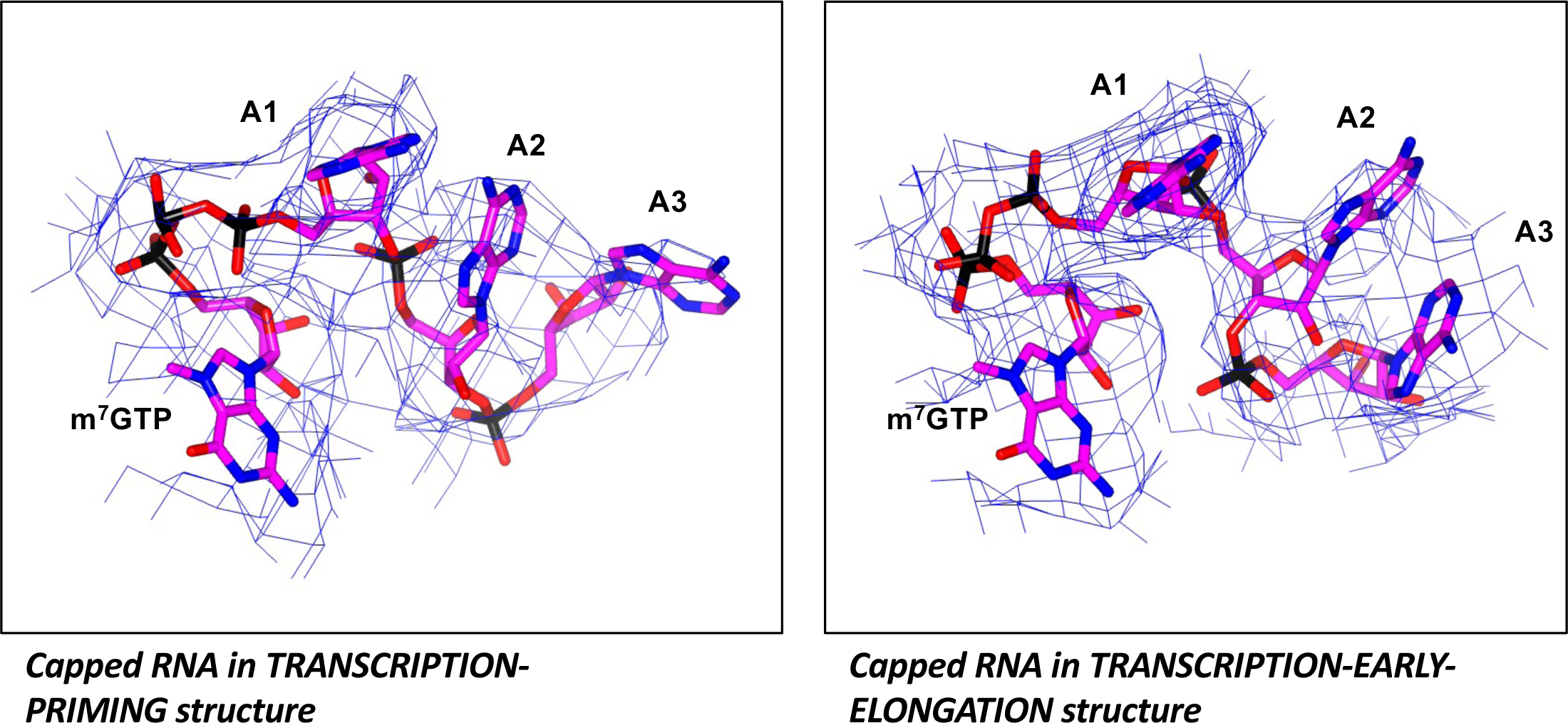
Quality of the map for the bound capped RNA. Zoomed-in views of the capped RNA (shown as sticks in magenta) in the **TRANSCRIPTION-PRIMING** and **TRANSCRIPTION-EARLY-ELONGATION** structures are shown with the relevant map overlaid. In both cases, the map is clipped to the capped RNA.

**Supplementary Figure 6.**
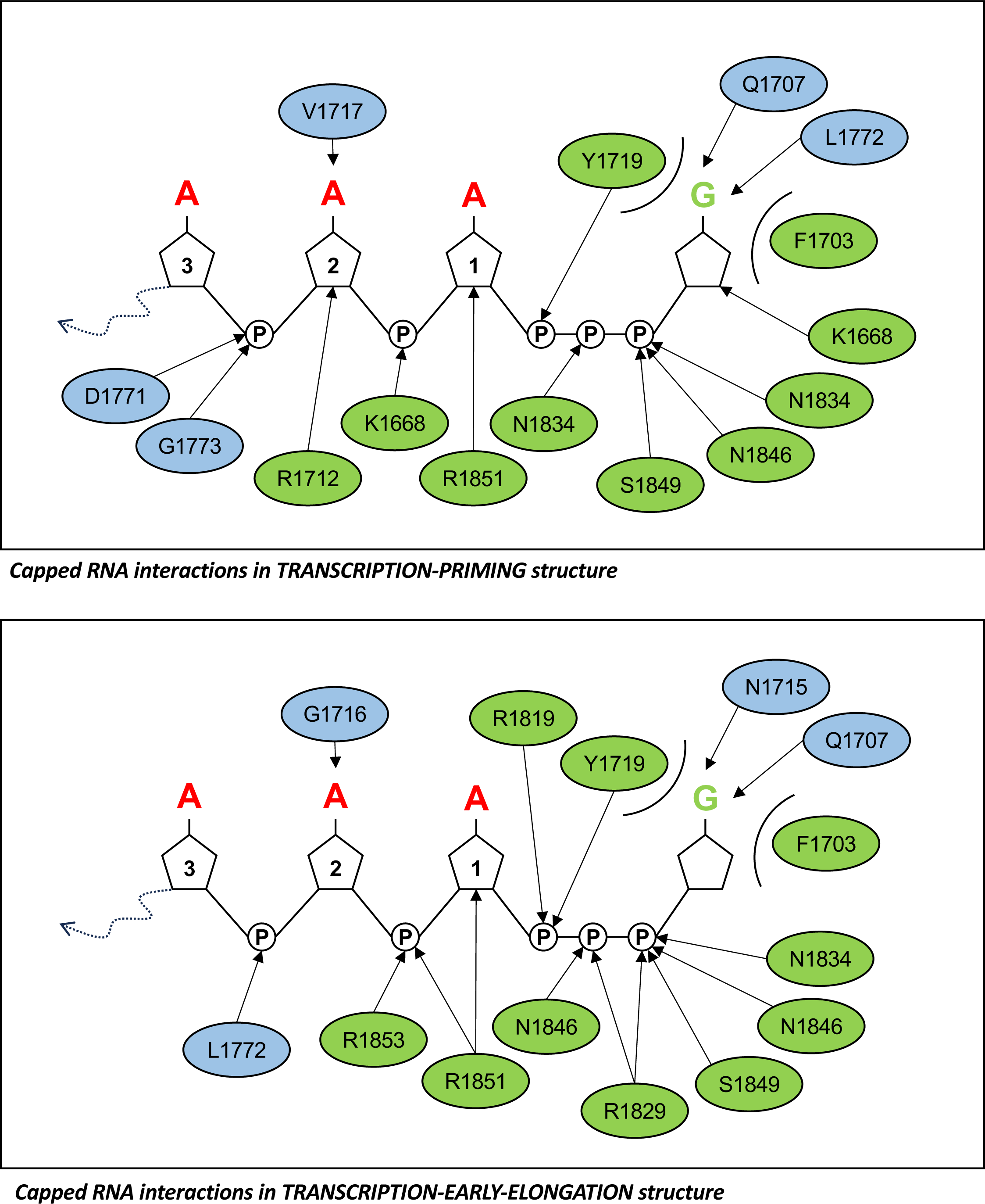
Capped RNA interaction map. A schematic overview of all interactions between SFTSV L protein residues and bound capped RNA is shown. Mainchain interactors are coloured blue, whereas sidechain interactors are coloured green.

**Supplementary Figure 7.**
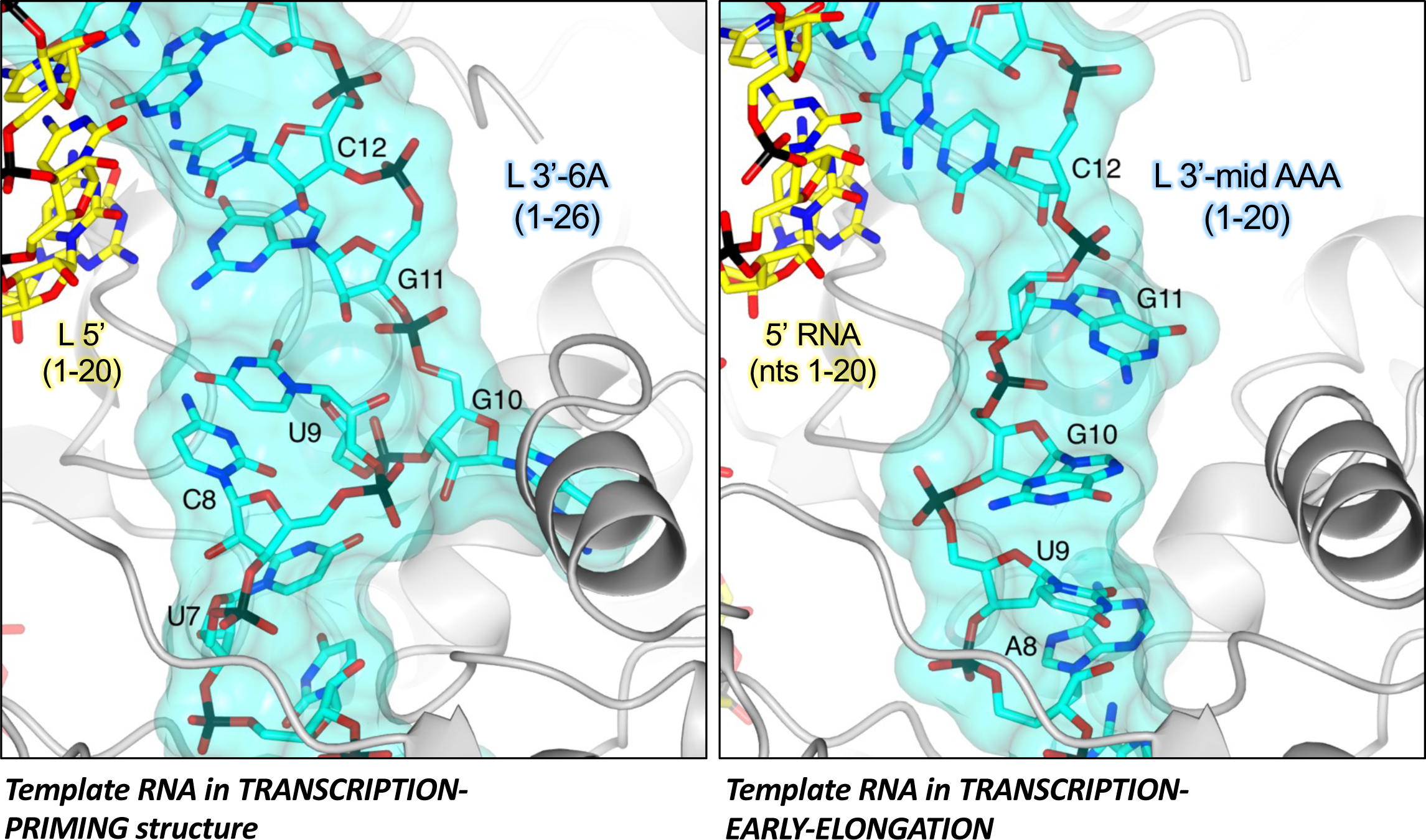
Status of the 3’ RNA between the distal duplex and the template entry channel. A zoomed-in view of the SFTSV L protein TRANSCRIPTION-PRIMING (left) and TRANSCRIPTION-EARLY-ELONGATION (right) are shown. RNA in both structures is shown as sticks with surface overlaid (30% transparency) and coloured either yellow (5’ RNA) or cyan (3’ RNA).

**Supplementary Figure 8.**
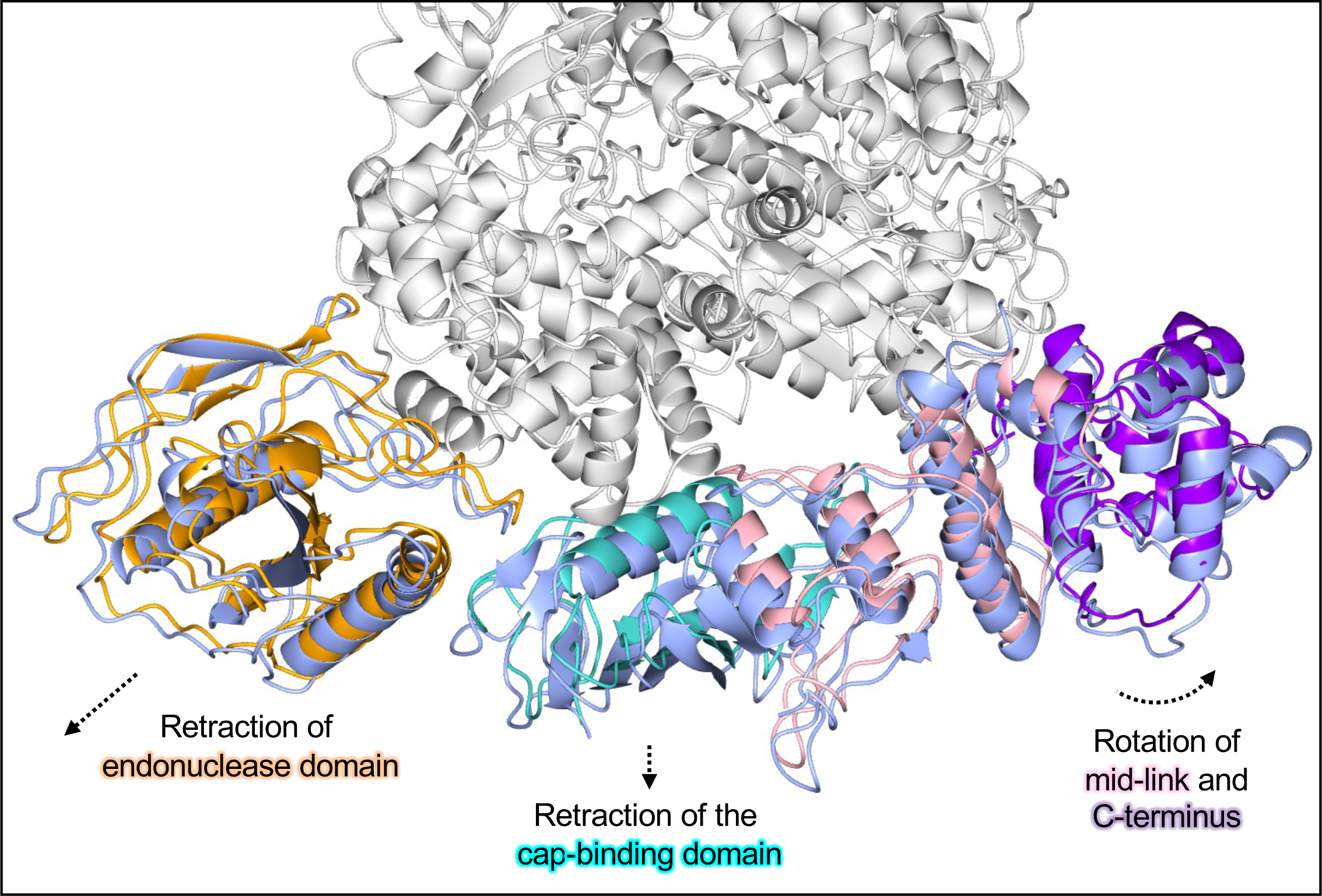
Conformational changes as the SFTSV L protein moves from transcription priming to a transcription-specific early elongation state. The SFTSV L protein **TRANSCRIPTION-PRIMING** and **TRANSCRIPTION-EARLY-ELONGATION** structures are shown. Key L protein domains in the TRANSCRIPTION-PRIMING structure are coloured canonically, i.e.: endonuclease (orange), cap-binding domain (cyan), mid-link domain (light pink), and extreme C-terminus (purple). For comparison, the same domains in the **TRANSCRIPTION-EARLY-ELONGATION** are each coloured ice blue. Dashed arrows indicate the direction of movement for each domain.

**Supplementary Figure 9.**
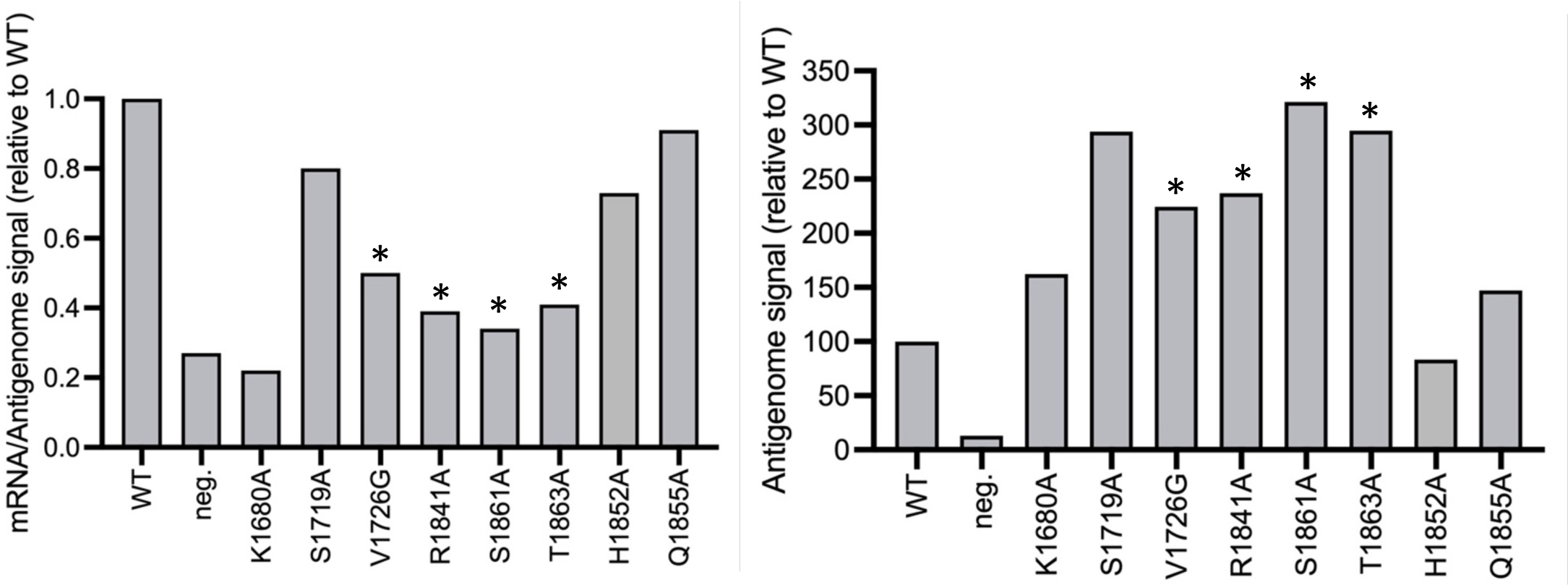
Northern blot quantification. (Left) RNA signals on northern blots for selected mutants were quantified using ImageJ/FIJI and the mRNA-to-antigenome signal ratio was calculated. (Right) Antigenome signals for selected mutants in northern blots were quantified via intensity profiles using ImageJ/FIJI. For both analyses, the wild-type ratio was set at 100% for each experiment allowing the comparison of independent experiments. Asterisks indicate mutants with a transcription-specific reduction. Data plotted in Prism.

**Supplementary Figure 10.**
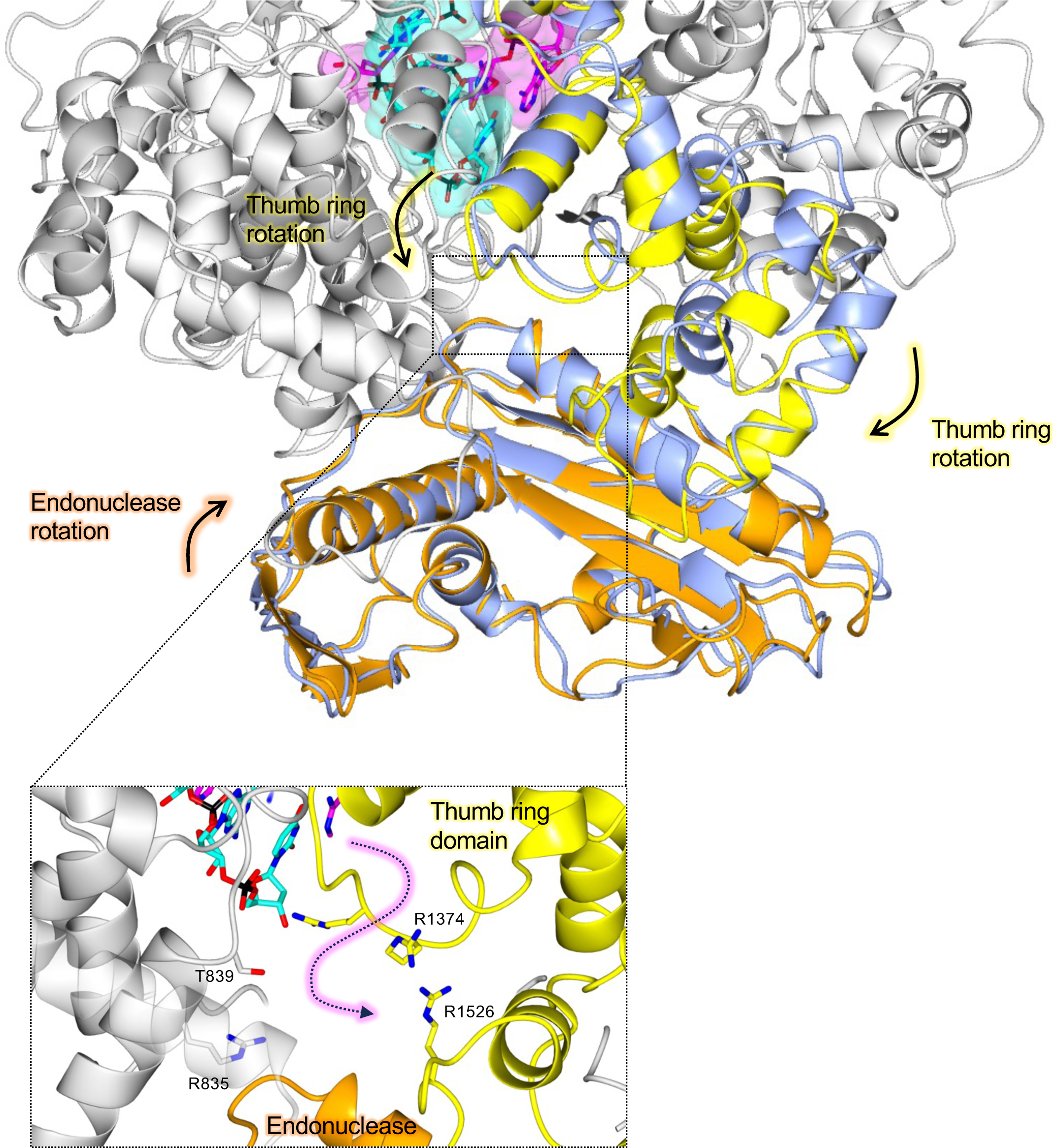
Insights into cap-independent primer-elongation. The SFTSV L protein **TRANSCRIPTION-PRIMING (*in vitro*)** structure is shown with the published apo L protein structure (PDB: 6Y6K) superposed by SSM in CCP4mg. Parts in grey are similar in both structures and, where this is true, only the **TRANSCRIPTION-PRIMING (*in vitro*)** structure is depicted. Domains of the **TRANSCRIPTION-PRIMING (*in vitro*)** structure that are shown to either rotate or remodel to enable primer entry and base pairing with the 3’ RNA are coloured (orange for the endonuclease and yellow for the thumb ring domain). The corresponding apo structure domains are shown in ice blue. RNAs are shown as sticks with surface overlaid (30% transparency) and coloured either cyan (3′ RNA) or magenta (primer RNA). Movements to the endonuclease and thumb ring domains precipitated by the progression to late elongation are indicated. Additionally, a zoomed-in view of the proposed product exit channel through which the primer has likely entered the RdRp core is shown. The path of the primer RNA, which is coloured magenta, is indicated by a dashed arrow.

**Supplementary Figure 11.**
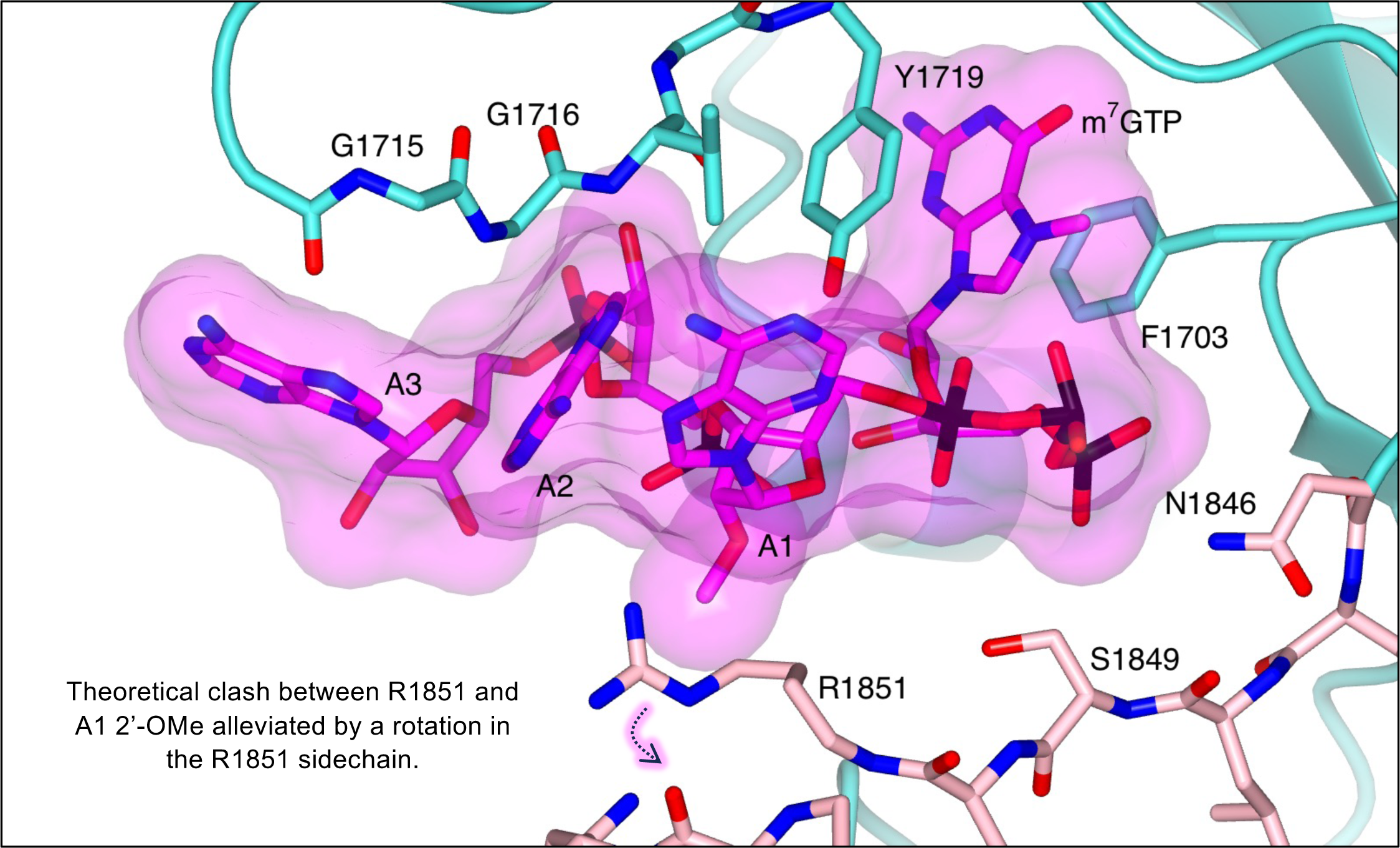
Exploring the effect of cap1 methylation. The cap-binding domain is shown with bound capped RNA. The cap-binding domain is coloured cyan, whereas the neighbouring mid-link domain is shown in light pink. The bound capped RNA is shown as sticks with surface overlaid (30% transparency). The primer RNA has been modified by the substitution of A1 with 2’-O-methyladenosine (PDB: A2M). Sidechains of key interacting amino acids, coloured according to the protein domain, are shown as sticks and labelled accordingly. The addition of the methyl group at the O2’ position would clash with the current placement of the R1851 sidechain, however, as indicated, the R1851 sidechain. could rotate and in doing so create sufficient space for cap1 RNA.

